# An experimental test of chronic traffic noise exposure on parental behaviour and reproduction in zebra finches

**DOI:** 10.1101/2021.11.30.470558

**Authors:** Quanxiao Liu, Esther Gelok, Kiki Fontein, Hans Slabbekoorn, Katharina Riebel

## Abstract

Chronic traffic noise is increasingly recognised as a potential hazard to wildlife. Several songbird species have been shown to breed poorly in traffic noise exposed habitats. However, identifying whether noise is causal in this requires experimental approaches. We tested whether experimental exposure to chronic traffic noise affected parental behaviour and reproductive success in zebra finches (*Taeniopygia guttata*). In a counterbalanced repeated-measures design, breeding pairs were exposed to continuous playback of one of two types of highway noise previously shown to be either neutral (control) or aversive. Parental nest attendance positively correlated with feeding effort and was higher for the aversive than the control sound and this effect was more pronounced for parents attending larger broods. However, neither noise condition affected offspring number, growth or body mass. The absence of an effect held when we combined our data with data from two other comparable studies into a meta-analysis. We discuss whether the increased nest attendance could be a compensatory strategy that alleviated detrimental noise effects on the chicks, and whether it could be caused by impaired parent-offspring or within-pair communication. Future work should test these hypotheses and investigate potential long-term costs of increased parental engagement.

## INTRODUCTION

Wildlife diversity and abundance decline near roads that expose animals to altered habitats, traffic hazard, chemical and noise pollution (Benítez-López et al., 2010; Newport et al., 2014; Bennett, 2017; Kunc and Schmidt, 2019). This negative association is especially well documented in birds (Reijnen et al., 1996; Bayne et al., 2008; Arévalo and Newhard, 2011; Herrera-Montes and Aide, 2011). Field data comparing noisy versus quiet sites within the same populations found, for example, noisy territories to be associated with lower pairing success in ovenbirds (*Seiurus aurocapilla*, Habib et al., 2007), lower hatching success in western bluebirds (*Sialia mexicana*, Kleist et al., 2018), smaller brood sizes in great tits (*Parus major*, Halfwerk, et al., 2011a) and eastern bluebirds (*Sialia sialis*, Kight et al., 2012) and reduced feeding and nestling growth in house sparrows *(Passer domesticus*, Schroeder et al., 2012).

These observations are suggestive of a causal relation between elevated noise levels and reduced breeding performance. However, the presence of roads also alters vegetation and food availability, increases chemical pollution and the danger of physical impact of moving vehicles. Any of these factors directly alters habitat quality in ways that could also affect breeding success. A number of experimental noise exposure studies addressed these potential confounds by breaking correlations between parental and territory quality and noise by cross-fostering chicks or by experimentally altering noise levels in the field (Table 1). Such noise treatments were found to affect a number of physiological and behavioural parameters, for example leading to a suppressed immediate corticosterone response in breeding female tree swallows (*Tachycineta bicolor*, Injaian et al., 2018a), altered territory choices in great tits (Halfwerk et al., 2016) and tree swallows (Injaian et al., 2018b), increased vigilance in house sparrows (Meilleŕe et al., 2015a), altered intra-pair communication in great tits (Halfwerk et al., 2011b) or parent-offspring interactions in tree swallows (Leonard and Horn, 2012; Leonard et al., 2015).

**Table 1.**
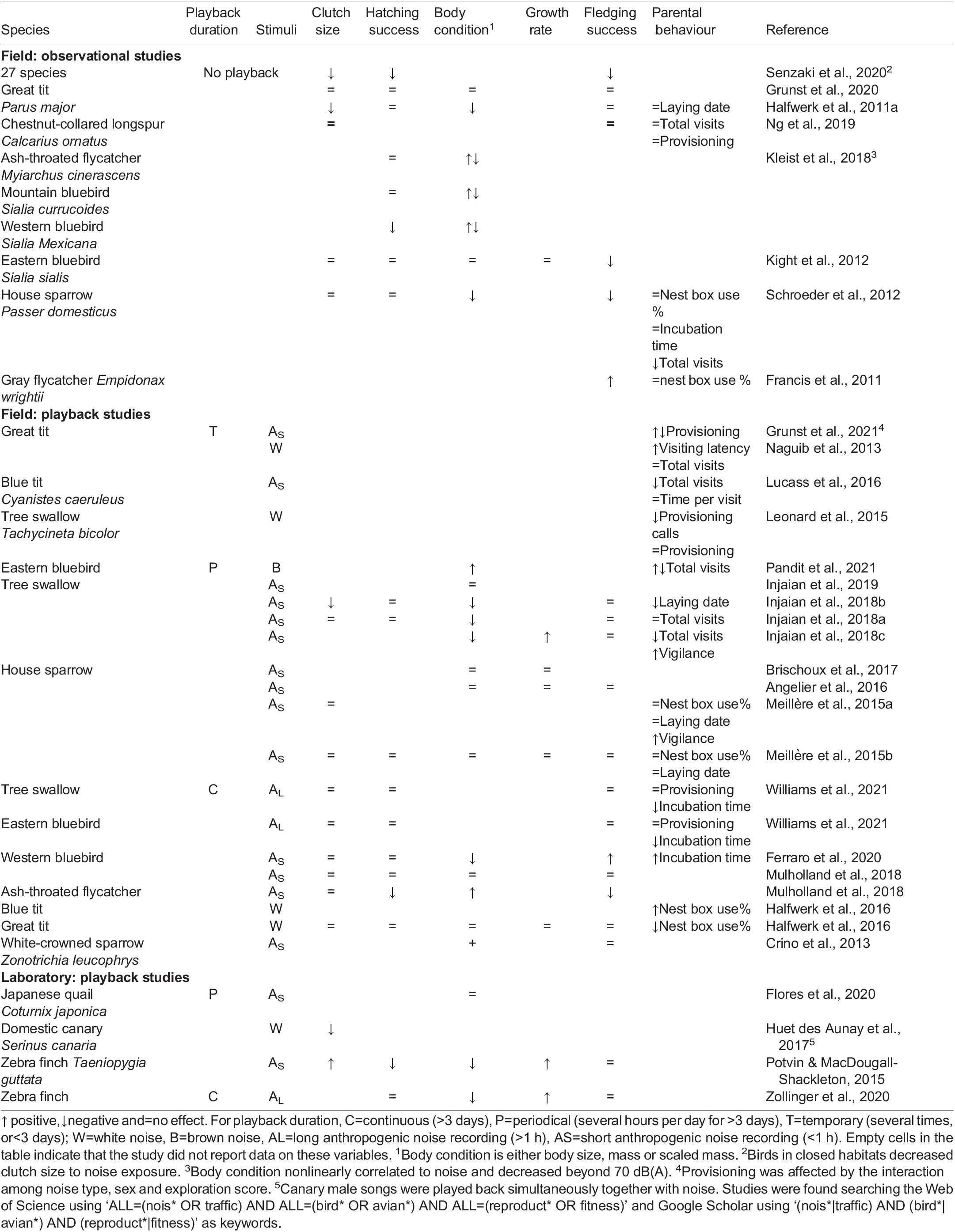
Studies investigating effects of anthropogenic noise on avian reproduction.

These observed physiological and behavioural effects of noise exposure, surprisingly, are not associated with a clear pattern of knock-on effects on reproductive outcomes across studies (neither within nor between species). As listed in Table 1, some studies reported negative effects of noise exposure on clutch size (Injaian et al., 2018b), hatching and fledging success (Mulholland et al., 2018) and offspring body condition (Injaian et al., 2018a,b,c; Ferraro et al., 2020), while others reported an absence of effects (Meilleŕe et al., 2015b; Angelier et al., 2016; Halfwerk et al., 2016; Injaian et al., 2019; Williams et al., 2021) or even mixed effects (Ferraro et al., 2020; Pandit et al., 2021) for some of the same measurements. The overall picture suggests a slew of physiological and behavioural parameters to change if birds are exposed to anthropogenic noise, but rarely, if ever, an effect on the net outcome of reproduction during the experimental period. However, we should be careful to draw any final conclusion from this set of first-generation studies, often only investigating one breeding event and varying in methodology, type of noise stimuli, and study species, as they also revealed how difficult it is to manipulate noise condition as a single parameter in the field (Table 1).

Added complications arise from the interactions between noise playbacks and other ecological factors, as noise playbacks in the field can affect organisms other than the study species. For example, noise can reduce arthropod abundance (Bunkley et al., 2017), which may contribute to habitat degradation for breeding birds. Species interactions could thus amplify, mask or confound potential effects of noise on breeding performance. For example, at and around North American gas extraction sites, avian predators avoided the noisiest areas near the sites, which is likely the reason for the observed improved breeding performance of their prey species at noisy sites compared to nearby control sites (Francis et al., 2009). These examples show that although many bird species avoid noise (McClure et al., 2013; Liu et al., 2020) others might not (Francis et al., 2009) and depending on context and species, field playback experiments could selectively sample bird species or individuals with higher noise tolerance and associated poorer breeding quality inadvertently (Naguib et al., 2013; Patrick and Weimerskirch, 2014). Sampling can also be biased, if lower quality individuals are more likely to settle for breeding in the poorer quality habitats near roads (Injaian et al., 2018b).

In contrast to field studies, playback experiments in the laboratory allow researchers to keep environmental factors constant which allows to separate direct versus indirect effects of noise on reproductive success. Huet des Aunay et al. (2017) tested whether noise affected female investment (clutch size) in domestic canaries (Serinus canaria), a species where male song is sufficient to stimulate egg laying even in the absence of a male (Kroodsma, 1976). Tested females were exposed to playbacks of male song and urban noise in two conditions: the noise was played in between or during songs, therefore exposing females to the same amount of overlapping (masking parts of the song spectrum) or nonoverlapping noise. In comparison with the non-overlapping, the overlapping noise treatment decreased clutch sizes, suggesting that masking noise interfered with the stimulating function of male song on the female’s reproductive system. Huet des Aunay et al. (2017) did not directly measure the associated stress physiological changes, but other studies have shown that urban or traffic noise exposure can lead to, e.g. increased levels of antioxidant glutathione (a potential marker of oxidative stress) in Japanese quail chicks (Flores et al., 2020), corticoid responses in tree swallows (Injaian et al., 2018a) and western bluebirds (Kleist et al., 2018), suppressed immune response to phytohemagglutinin in zebra finches (Brumm et al., 2021), and shortened telomeres in tree swallows (Injaian et al., 2019), great tits (Grunst et al., 2020), house sparrows (Meillére et al., 2015b) and zebra finches (Dorado-Correa et al., 2018).

Notably, most studies to date focused on the effects of prolonged or chronic noise exposure on either parents or offspring, but rarely on both parents and offspring. Two such studies (Potvin and MacDougall-Shackleton, 2015; Zollinger et al., 2020), however, have been conducted in zebra finches, an important model for avian development (Griffith and Buchanan, 2010). Breeding pairs exposed to playbacks of urban park noise with edited-in vehicle sounds had an increased embryo mortality compared to a no-playback control group (Potvin and MacDougall-Shackleton, 2015). However, the overall breeding outcome (i.e. the number of fledglings) did not differ between the treatment and control group suggesting that the noise exposed birds initially had laid more eggs. A comparable study that also exposed breeding zebra finches either to traffic noise or to a control treatment without playback (Zollinger et al., 2020), found no treatment effects on the embryo mortality rate or the number of offspring. In both zebra finch studies (Potvin and MacDougall-Shackleton, 2015; Zollinger et al., 2020), the offspring mass at early age was, however, negatively affected by the noise treatment. Early mass is an important predictor of adult condition in zebra finches (Buchanan et al., 2004; Krause et al., 2017), thus showing that the noise treatment had an effect on offspring development. There were no behavioural observations in these studies, so it is not clear whether the noise had a direct effect on chick development or that the effect was indirectly arising from altered parental behaviour. Experimental studies monitoring both parental behaviour and chick development could help to answer the question of whether negative noise impact is directly acting on chicks or indirectly via their parents.

The field of experimental noise exposure studies in birds is growing quickly but also still developing its paradigms, as becomes apparent when comparing the type and duration of playbacks used across studies (see Table 1). Ideally, any experimental study should compare carefully selected noise stimuli to a suitable control treatment. Studies to date have tested a great array of different types of anthropogenic noise (or digitally synthesized forms of it) as stimuli but mostly compared a noise-exposed group to a control group not experiencing any playbacks. A no-playback group provides the clearest contrast to a playback group to demonstrate adverse effects of a noise stimulus. However, such a design cannot separate noise specific effects from effects of experiencing a sudden change from a quiet to a noisy environment or from effects arising from hearing unknown, novel sounds.

We therefore designed a study to compare two types of highway noise, for which we had previously tested that they were either behaviourally neutral or aversive to adult zebra finches from our colony (Liu et al., 2020). Briefly, for these earlier tests, small groups of zebra finches had been temporarily housed in two interconnected aviaries that differed in the presence and/or intensity of traffic noise playback. The birds could fly freely to express spatial preferences with respect to sound exposure levels. For the breeding experiment reported here, all birds had been previously tested in this setup with all combinations of quiet (no noise), moderate- and high-intensity noise. The tested birds preferred for quiet aviary over high (but not moderate) intensity traffic noise. Based on these previous tests (Liu et al., 2020), we selected one 24-h-long continuous recording of each category of traffic noise, the aversive (high-intensity) traffic sound level as experimental treatment and the behaviourally neutral (not avoided, moderate-intensity) traffic sound level as control noise treatment for the current breeding experiment. In a fully balanced design with cross-over (Fig. 1), each pair was allowed to breed twice: once while exposed to the aversive and once to the control traffic noise recordings. By simultaneously monitoring parental behaviour and reproductive outcomes of the same breeding pair under two types of noise playbacks, we aimed to investigate whether potential detrimental effects on chicks were co-occurring with altered parental behaviour or whether the noise altered parental behaviour but not the chicks’ development and survival. Based on earlier reports on the negative impact on reproductive output and offspring physiology in zebra finches (Potvin and MacDougall-Shackleton, 2015; Dorado-Correa et al., 2018; Zollinger et al., 2020), we expected to find a negative effect of the aversive noise on offspring development. If these changes are (partly) driven by changes in parental behaviour both nest attendance and offspring body mass should be affected. Adding observations of parental behaviour to chicks’ biometry markers of development, should help revealing associations between the two, thus helping to develop further regarding the underlying mechanism of anthropogenic noise on avian reproduction. We also conducted a post-hoc meta-analysis combining our data with two other published noise exposure breeding studies (Potvin and MacDougall-Shackleton, 2015; Zollinger et al., 2020) to examine the overall effect size of noisy rearing conditions on zebra finch reproduction.

**Fig. 1.**
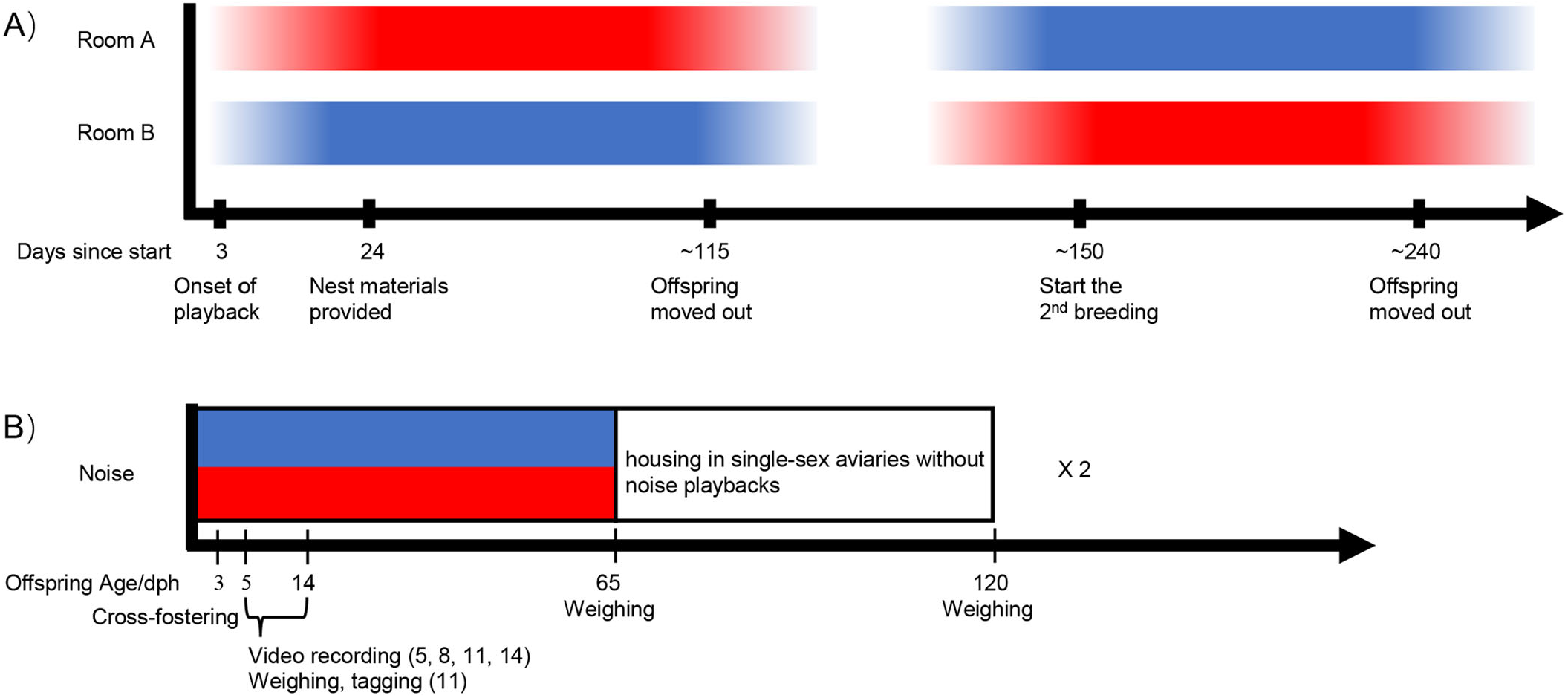
Timeline and design of the repeated measures breeding experiment (A) and the timeline for the experimental procedures within offspring cohorts (B). Noise playback type is indicated by colour (red=aversive, blue=control) and the colour gradient indicates the relative intensity of the noise playback compared to the peak level. The x-axis in panel A shows the number of days since the start of the experiment. In each breeding round, the noise playback faded in 2 weeks before nesting materials were provided to the breeding pairs and the playback lasted until offspring from all breeding pairs reached 65 days old. The x-axis in panel B shows offspring age (in days post hatching) in relation to the reported procedures.

## RESULTS

### Reproductive outcomes

The females of the majority of pairs (23/30) laid eggs in both breeding rounds (for details, see Table S1). Females of six pairs only laid eggs in one breeding round (four in the control and two in the aversive noise condition) and one female did not lay eggs in either condition. Among pairs that had eggs, 13 pairs successfully hatched offspring in both breeding rounds, but a number of pairs that had clutches had no hatchlings: six pairs in the control noise, four pairs in the aversive noise and another four pairs in both conditions. The latency to the first egg, clutch size, the number of successfully hatched chicks and the number of unhatched eggs did not differ between the two noise treatments, breeding rounds or the order of exposure to the two different noise treatments (see Fig. 2A–D, Table 2, for rejected interactions, see Table S2). In addition, the number of successfully hatched chicks per pair did not differ from that of normally bred birds in our colony (see Table S5). Clutch weight (only measured in the first round, see ‘Materials and Methods’) was not affected by noise playback treatment, and this held both for the sample including or excluding the pairs that had not produced a clutch (see Table S6). Likewise, total hatchling weight per brood (only measured in the second round, see ‘Materials and Methods’) was not affected by playback type including or excluding the pairs that had not laid any eggs (see Table S6).

**Table 2.** Generalised linear mixed model analyses of breeding outcomes across the two noise treatment groups: (a) latency to the first egg, (b) clutch size, (c) successfully hatched chicks and (d) number of unhatched eggs.

| Effects | $\chi^2$ | P-value |
| --- | --- | --- |
| a) Latency to the first egg |  |  |
| Noise | 0.02 | 0.89 |
| Round | 2.00 | 0.16 |
| b) Clutch size |  |  |
| Noise | 0.10 | 0.81 |
| Round | 1.00 | 0.32 |
| c) Successfully hatched chicks |  |  |
| Noise | 1.25 | 0.26 |
| Round | 1.25 | 0.26 |
| d) Number of unhatched eggs |  |  |
| Noise | 0.62 | 0.43 |
| Round | 0.08 | 0.78 |

**Fig. 2.**
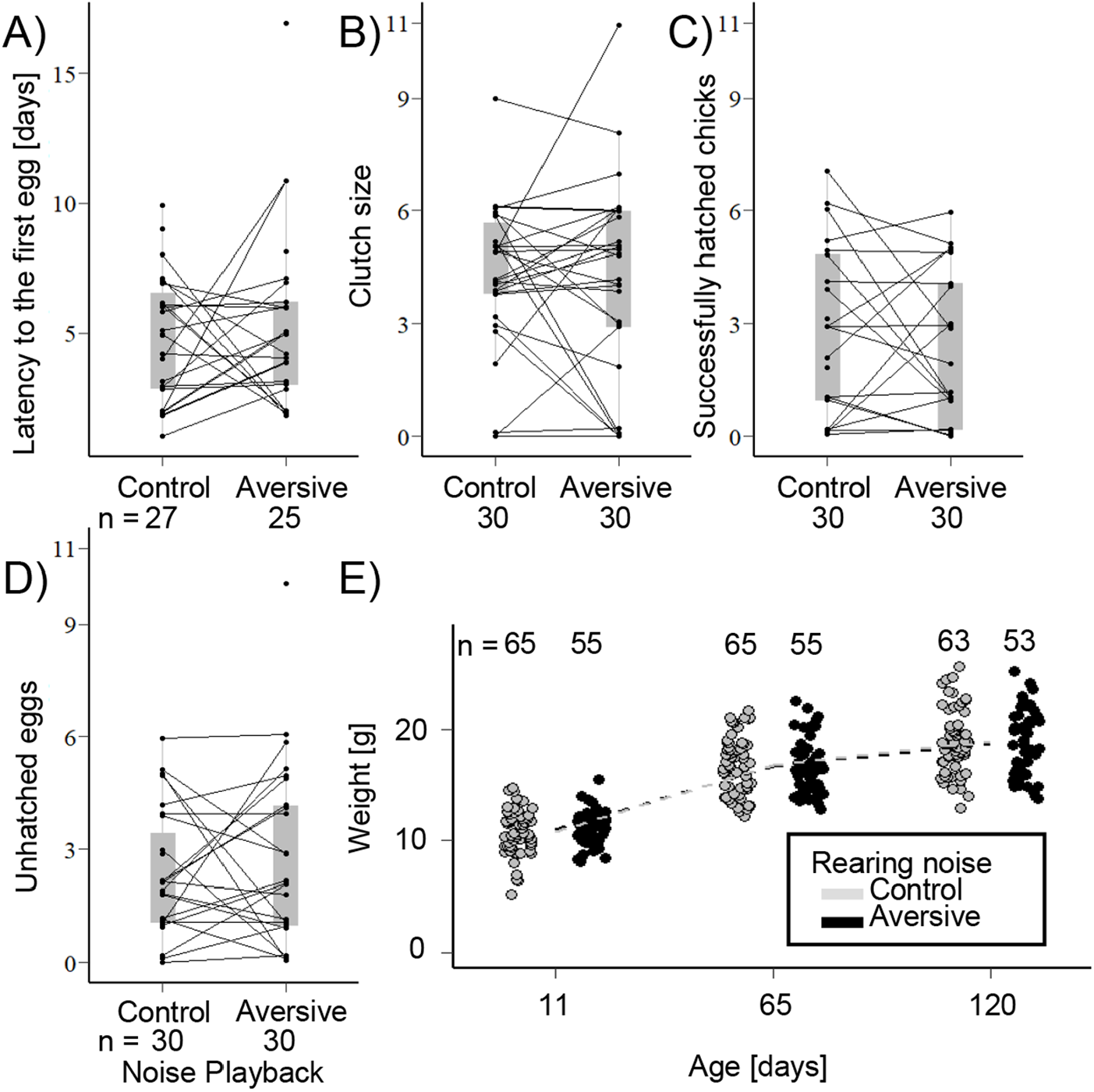
Reproductive performance during continuous exposure to two different levels of traffic noise. (A) Latency to the first egg,(B) clutch size, (C) successfully hatched chicks,(D) number of unhatched eggs. Each dot represents one pair during one breeding event (statistical details see Table 2). Two dots connected by lines belong to the same pair and show the two values obtained in each of the two breeding rounds. Grey overlayed boxplots give the median, 1st and 3rd quartile per treatment group. (E) Offspring mass at 11, 65 and 120 days. Each dot represents a single bird (statistical details see Table S3). For visualisation, overlapping data were randomly and horizontally jittered. Details on individual pairs’ breeding outcomes can be found in Table S1.

Offspring mass at the age of 11, 65 and 120 days old decreased with brood size but not in relation to noise playback type (see Fig. 2E, Table S3) and there were no significant interactions between these terms.

### Meta-analysis: effect of sound exposure on zebra finch breeding outcomes

To date, the present and two other studies (Potvin and MacDougall-Shackleton, 2015; Zollinger et al., 2020) have investigated the effect of noise exposure on breeding success in zebra finches. Combining the results in one meta-analysis shows no overall effect of chronic exposure to anthropogenic noise on hatching failure (effect size [95% CI]=−0.24 [−2.75, 2.27]) in zebra finches (Fig. 3A). Similarly, there was no effect of noise exposure on offspring mass (Fig. 3B): at the combined developmental stages (−0.17 [−0.40, 0.07]) or any of developmental stages when analysed separately: hatchling (−0.39 [−1.18, 0.40]), fledgling (−0.04 [−0.53, 0.45]) subadult and adult (−0.09 [−0.42, 0.34]).

**Fig. 3.**
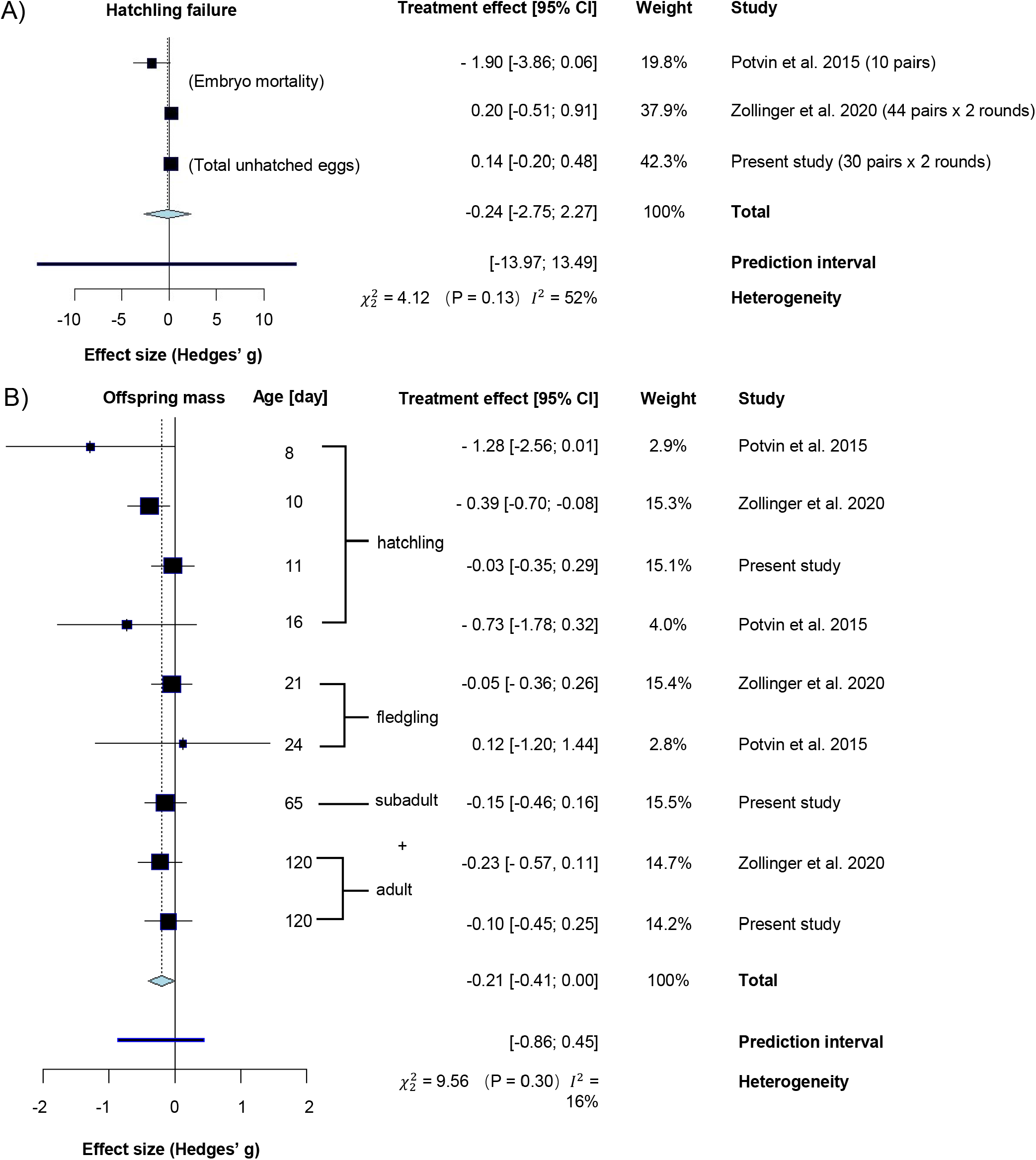
Forest plot showing the effect sizes from a meta-analysis testing the effect of traffic noise exposure on (A) hatching failure and (B) offspring mass. Squares in the forest plot give the treatment effect with 95% confidence intervals. Negative values mean negative effects from noise exposure. The size of the squares reflects the weight of the data in the meta-analysis. The dashed line indicates the total treatment effect.

### Parental behaviour

Males and females spent more time attending the nest when exposed to the aversive noise than to control noise across all measurement moments (median brood age 5, 8, 11, 14 days; effect size [95% CI]=0.40 [0.19, 0.62]). Nest attendance was overall negatively associated with brood size and brood median age. Males spent overall less time attending nests than females (Table 3a). Following analyses that combined data from both sexes of the same pair showed the same result in terms of effects of noise, brood size and brood median age (Table 3b, Fig. 4). However, noise exposure did not affect the overall number of combined nest visits (Table 3c) during the recorded period. The rejected interaction terms from aforementioned models can be found in Table S4.

**Table 3.**
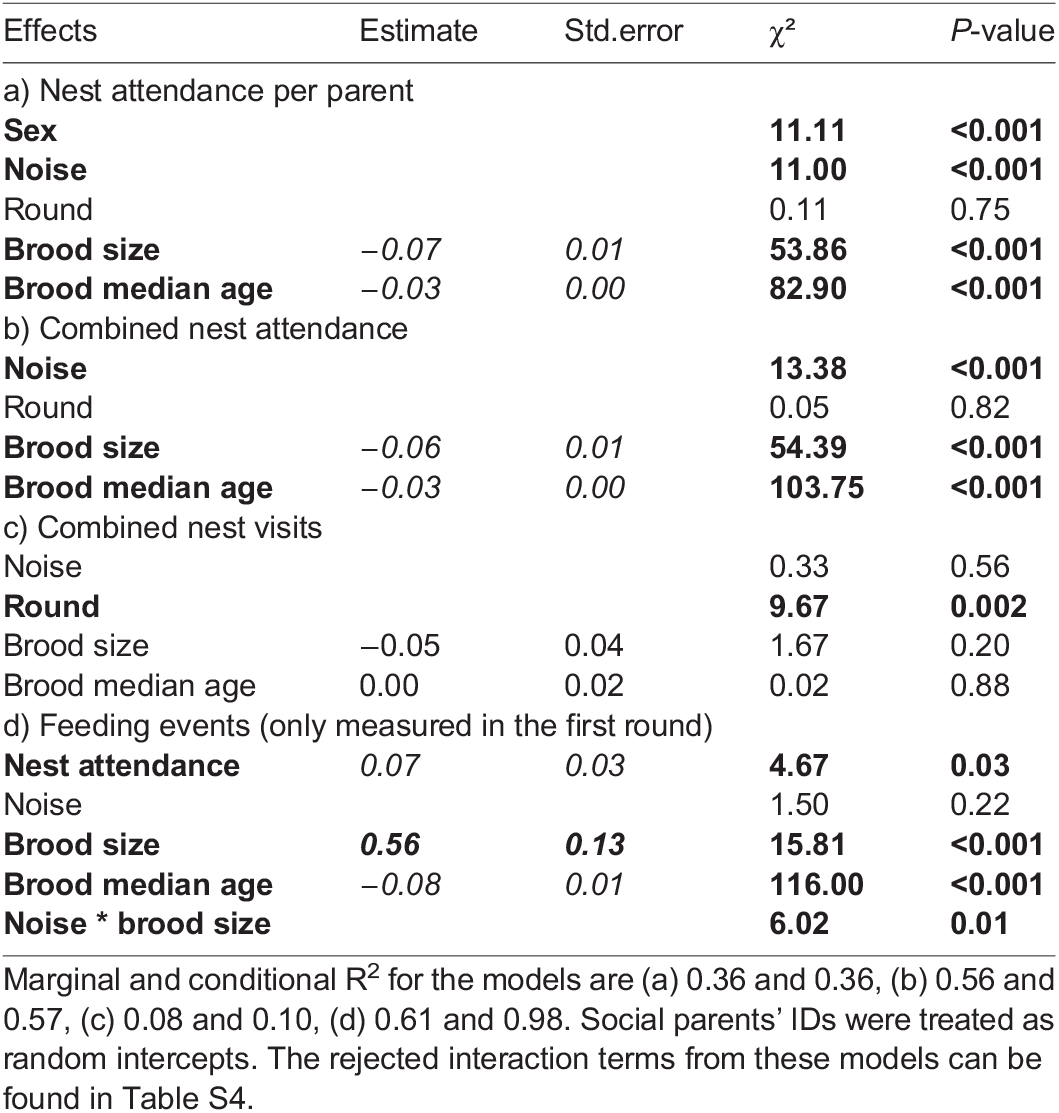
Testing for effects of the two types of noise playbacks on parental behaviour with generalised linear mixed model analyses.

**Fig. 4.**
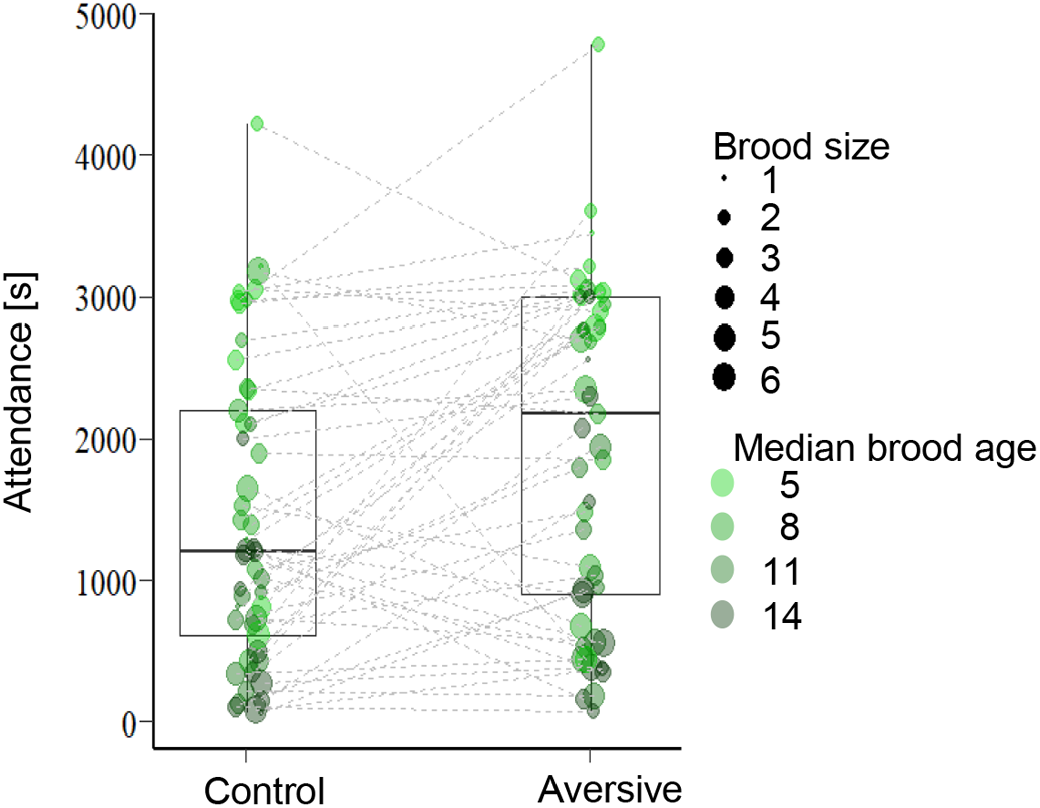
Parents’ nest attendance during chronic noise exposure. Each dot shows the combined time of both parents spent in the nest during a 50 min continuous sampling period during four different median brood ages (median brood age=5, 8, 11, 14, data extracted by blinded video analyses). Each pair was tested in both conditions and connected dots show two data points of the same parents attending their broods at the same median age during control or aversive noise playback. The size and colour saturation of the dots code for the brood size and median brood age. Dots that are connected by lines show the same breeding pair’s behaviour during either the aversive or control noise exposure at the same median brood age. Data are from *N*=2×49 median-brood-age paired videos from 13 pairs that had chicks in both breeding rounds.

The in-nest video recordings during the first breeding round showed that combined nest attendance of both parents predicted the total feeding events (Fig. 5, Table 3d). Besides, the parents fed the chicks more often if the brood was younger. The significant interaction between brood size and playback type showed that when exposed to the aversive noise, the parents with larger brood increased their feeding rate more than the parents with smaller brood (Fig. 5, Table 3d).

**Fig. 5.**
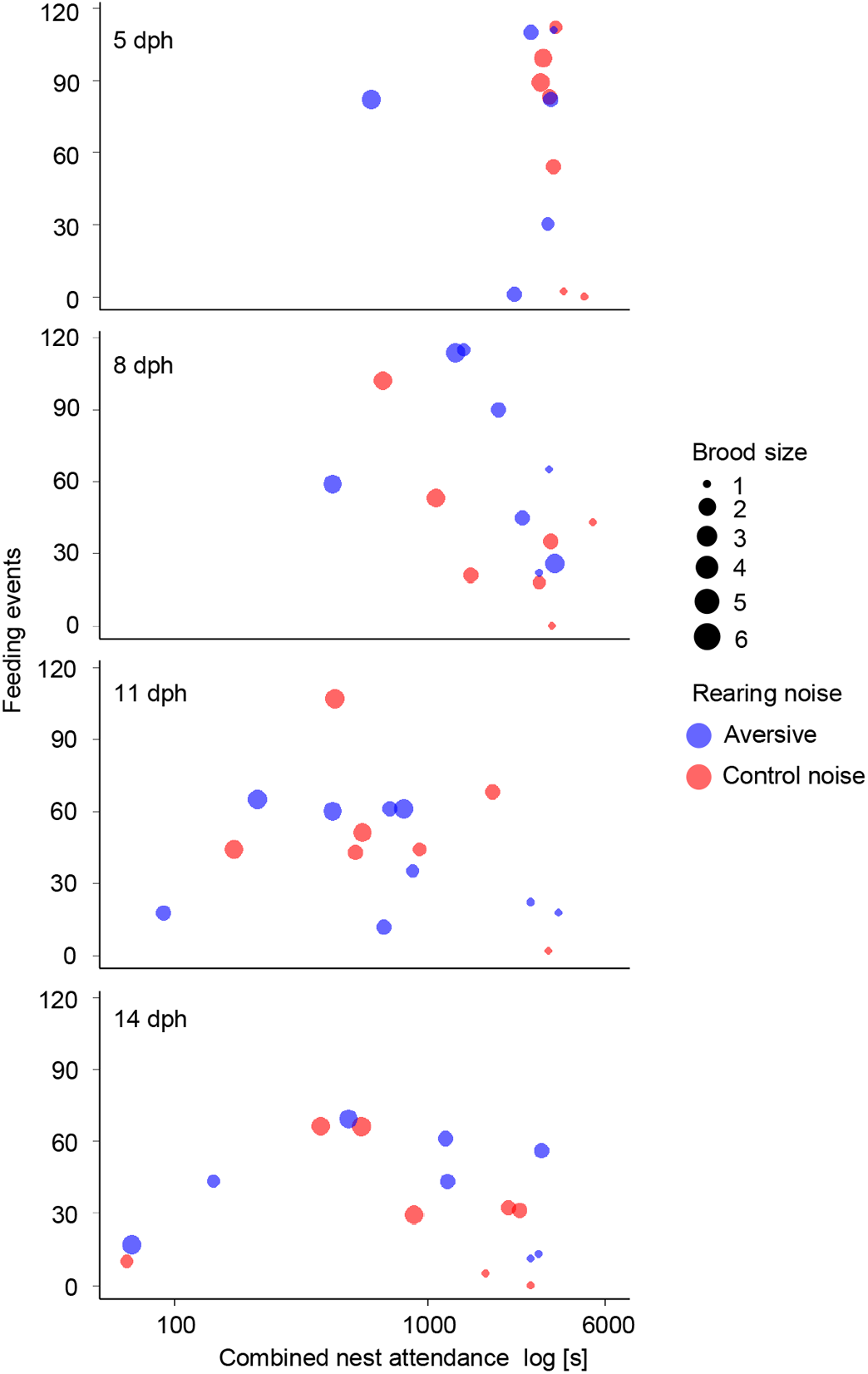
Combined nest attendance (the total time spent by the male and female in the nest) in relation to the number of feeding events. The x-axis is plotted on a logarithmic scale. Each row shows the data from each recorded median brood age. Each symbol represents the data from one video recording (*N*=66) of one pair during the first breeding round exposed to either control (blue) or aversive (red) noise. Dot size increases with brood size (1-6).

## DISCUSSION

The different traffic noise exposure conditions in our study affected parental behaviour, but the type of noise treatment had no significant impact on reproductive output. In both breeding rounds, parents spent more time attending the nest when exposed to the high-intensity, aversive noise compared to moderate-intensity control noise. Feeding rate, which was positively correlated with nest attendance, was higher in larger broods and more so if parents were exposed to the aversive than the control noise. While the parental behaviour changed according to noise treatment in both replicate groups there were no significant effects of the noise treatment on any of the measurements on chick growth or survival. There was no effect from the noise treatments on chick mass, but in line with previous reports in this species, chick mass was affected by brood size. Our meta-analysis of the combined data sets of our and two other similar zebra finch exposure studies, showed no overall effect from traffic noise exposure on hatching failure or offspring mass.

The impacts of noise exposure on parental behaviour are in line with other studies reporting behavioural effects of noise on zebra finches fitness-relevant contexts, such as mate choice, intra-pair communication, vigilance, foraging and learning about new food sources (Swaddle and Page, 2007; Villain et al., 2016; Evans et al., 2018; Liu et al., 2020; Corbani et al., 2021; Osbrink et al., 2021). Against this backdrop, the absence of an effect on reproductive success and offspring condition seems at first surprising and also to differ from the findings of an earlier, comparable noise exposure study in zebra finches (Potvin and MacDougall-Shackleton, 2015) where traffic noise exposure led to more nesting attempts, delayed egg laying dates, and a higher embryo mortality rate than in pairs in a control group breeding without noise exposure. However, our meta-analysis, combining the data of all three noise exposure studies in zebra finches to date, showed no overall effect from traffic noise exposure on embryo mortality or offspring mass. This suggests that the findings of the individual studies that differed in the direction or presence of effects might have arisen from low power of the individual studies (see the ‘weight’ in Fig. 4). This interpretation seems more likely than that the differences were caused by the different playback protocols, as our data (two types of noise) and the data from one of the two studies using a noise versus silence treatment (Zollinger et al., 2020) both showed an absence of effects on measures of reproductive failure. It is worth stressing that there are methodological differences across the three studies on the noise effects in zebra finches to date, concerning the type of noise, the exposure scheme, at what age the biometric measures were taken, whether the control treatment consisted of no noise or low-level control noise and whether a single measure or repeated measures design was used. Moreover, a cross-fostering paradigm like used in our study could mask or amplify the impacts of noise on breeding success because parents were not choosing their brood size. However, our results regarding an absence of effect on reproductive efforts are consistent with the Zollinger et al. (2020) study, in which the offspring were not cross-fostered. Clearly more studies in this and other species are needed assessing not only single breeding events but lifetime reproductive success to reach more general conclusions regarding the impact of noise on reproduction.

In contrast to the absence of an effect on the reproductive output parameters the noise clearly affected the behaviour of the parents: the aversive noise exposure increased nest attendance overall (effect size [95% CI]=0.40 [0.19, 0.62]) and increased feeding rates of parents with a larger brood in particular. Unfortunately, there are no data on parental behaviour in the other two studies, but our findings are unlikely to be spurious as the birds were tested in a paired-measures design, and the change in nest attendance showed a substantial effect size. This suggests that, although the reproductive outcome was not affected, parents exposed to the aversive high-intensity noise might have put in extra efforts during chick rearing. It seems that (at least in the relatively benign environment of the laboratory), parents might be able to buffer some direct negative impacts of noise. This observation should be replicated first in other studies/populations, but such an interpretation is in line with the absence of an effect on chick biometrical measures in our study or the observations that initial lower weight in noise exposed chicks disappeared later on in the other two experimental noise exposure studies in this species (Potvin and MacDougall-Shackleton, 2015; Zollinger et al., 2020).

The increased time and feeding investment adds to a list of other behavioural effects of noise on zebra finches: pair bond erosion (Swaddle and Page, 2007), intrapair communication changes (Villain et al., 2016), increased vigilance (Evans et al., 2018) and decreased cognitive performance (Osbrink et al., 2021). Moreover, some effects might not show up immediately but only later in life, such as the reduction in telomere length observed in offspring raised in high noise conditions (Dorado-Correa et al., 2018) of the aforementioned noise exposure study (Zollinger et al., 2020). The changes in parental behaviour we observed during high noise exposure could also entail potential long-term costs for zebra finches: higher effort foraging regimes affect lifetime expectancy in individuals that were of relatively lower condition as juveniles (Briga et al., 2017). Whether the increased parental nest attendance also carries such a hidden costs in lifetime expectancy and reduces lifelong reproduction needs to be addressed in future studies.

The absence of an effect on reproductive success in benign laboratory conditions that offer abundant food, near constant climate and no predation risk is not necessarily in contrast with negative effects on breeding behaviour reported in field studies on other species (Naguib, 2013; Leonard et al., 2015; Lucass et al., 2016). The general picture arising from these studies is an overall negative effect of high noise levels on both parental visit rates and offspring condition, likely due to interference by noise on parent-offspring (Leonard and Horn, 2012; Schroeder et al., 2012) or parent-parent communication (Villain et al., 2016). For example, young blue tits beg more during playback of heterospecific sound than during species-specific calls or silence (Lessells et al., 2011) and continuous white noise increases begging in tree swallows (Leonard et al., 2015), which could be an explanation for why the parents increased their nest attendance time. However, noise can also mask chick begging, so if parents do not hear begging calls well, or if chicks do not respond to parents arriving with food by begging (Leonard and Horn, 2012), feeding rates go down, which will cause a negative impact on chick growth. This will be much more pronounced in the field where parents have to work harder to find food, face additional impacts from roads (pollution) and increase their vigilance in a noisy habitat (McIntyre et al., 2014; Meilleŕe et al., 2015a). These complications could burden breeding parents and limit how much resource they would divert to mitigate the effects of noise on their chicks. Such differences between field and laboratory conditions and also between species could explain why we did not find the reduced nest visits that had been observed in house sparrows in the field (Schroeder et al., 2012). It may also be that masking effects on communication have a different, probably larger impact on birds in real-world situations, compared to zebra finches in laboratory housing in small cages, with close proximity between all birds.

Additionally, zebra finches are opportunistic breeders originating from semi-arid zones with unpredictable food supplies and they are quite resilient to environmental challenges like extreme temperatures and nutritional stress (reviewed by Griffith et al., 2021). For example, the average egg mass and fledgling rate in zebra finches are not easily affected by variation in quality of their food (Griffith et al., 2021). This suggests that zebra finches are perhaps also less susceptible to potential detrimental effects of chronic traffic noise exposure than other species. Seasonal breeders, for example, rely on other environmental and intrapair cues than lifelong opportunistic breeders. However, other aspects of zebra finch biology point to potential vulnerability to noise: synchronisation of breeding activity (Waas et al., 2005; Brandl et al., 2019) and parental communication (Villain et al., 2016) are all mediated by vocal communication (calls and singing), while zebra finch vocalisations carry only short distance compared to many other songbirds (Loning et al., 2021).

With so many effects on the behaviour of breeding birds of different species, the absence of short-term effects of experimental noise exposure experiments on reproductive success needs to be interpreted with caution: an absence of short-term effects is not predictive of potential long-term consequences (Santos and Nakagawa, 2012; Williams, 2018) and effects of chronic noise exposure might become apparent only later in life. For example, even in the absence of measurable differences in mass and in aspects of song performance known to be condition-dependent in noise-reared chicks when compared with control chicks at 120 dph (Zollinger et al., 2020; Liu et al., 2021), birds at the same age from these cohorts differed from control birds in telomere attrition rate, which has been linked to life expectancy (Dorado-Correa et al., 2018). We did not measure changes in physiology in either parents or offspring and we, therefore, cannot dismiss or exclude impacts that have been reported in quite a few studies (Injaian et al., 2018c; Kleist et al., 2018; Flores et al., 2020; Walthers and Barber, 2020; Zollinger et al., 2020).

In our study, reproductive success was unaffected, but birds attended their nests longer and fed their offspring more often in the high-intensity, aversive noise condition than in the control condition. These results are in line with the parental compensation hypothesis that posits that parents may flexibly adjust their parental care to more demanding circumstances and thus buffering negative environmental impacts (Rehling et al., 2012; Vitousek et al., 2017). Increased nest attendance (and feeding rate) may have compensated direct negative effects (see also Pandit et al., 2021 for a similar result in bluebirds). Leonard et al. (2015) also found that insect-eating tree swallow parents fed their chicks more often with elevated noise levels, although this appeared not sufficient to prevent a detrimental impact of noise on offspring body condition in this species (Injaian et al., 2018c). Detrimental effects of white noise on immune responses in tree swallows was only found among light nestlings in large broods (Obomsawin et al., 2021). This, like our results, suggests that large broods may be particularly vulnerable to noise impacts and parents might need extra parental care to compensate the detrimental effects of noise. Observations in other birds are similar to the findings we reported here: in white-crowned sparrows, for example, paternal nest attendance and feeding rate were found to be higher for nests closer to a road (Crino et al., 2011). Since our setting allowed us to exclude confounding factors like food availability and parental quality, the combined results from our study and other field studies (Injaian et al., 2018b; Pandit et al., 2021) support that traffic noise affects parents and that increased feeding rates could be one way to mitigate negative effects of noise. However, we do not know whether zebra finches can compensate the negative effects of noise in the long-term, or how such behaviour would affect the parents. Our study showed behavioural flexibility of breeding zebra finches with no apparent short-term consequences, but more experimental studies are needed on long-term fitness consequences of breeding in aversive noise.

Another, not mutually exclusive explanation for the increased nest attendance could be that the communication between the zebra finch parents was undermined by the noisy conditions. Zebra finch pairs use duet-like vocal exchanges to coordinate the parental behaviour (Boucaud et al., 2016, 2017) and the level of coordination predicts reproductive success (Mariette and Griffith, 2012, 2015). Noise from wind for example, has been shown to structurally change zebra finch duets and spatial proximity (Villain et al., 2016). It is therefore possible that traffic noise exposure in our experiment affected vocal communication and made the birds stay longer at the nest due to interrupted duty relief. Similarly, interrupted communication between parents and offspring as has been observed in tree swallows (Leonard et al., 2015) and blue tits (Lucass et al., 2016) may also occur in zebra finches and could have increased nest attendance time. Although zebra finch vocalizations are relatively soft compared to other songbirds (Loning et al., 2021), they should still be at least partly audible in the close proximity conditions in the small-cage environment of our laboratory experiment making masking of parental vocal communication perhaps a less likely explanation for the current observations. However, parents also react to the vocal signals of their chicks and the amount, timing and audibility of these signals might have been affected by the noise treatment in ways that made parents change their nest attendance behaviour.

In conclusion, we found further evidence for noise-related effects on zebra finch breeding behaviour. Aversive, high-intensity, traffic noise changed how often zebra finch parents fed and for how long they attended their nestlings, even under benign laboratory conditions. In line with other studies in zebra finches and a meta-analysis of these data, we observed no negative impact on reproductive outcomes during the experiment. Future studies are required to test whether this is due to the flexibility of this nomadic and opportunistic breeder or whether benign conditions in laboratory breeding colonies buffer noise impact, or whether there are long-term costs to the altered parental behaviour. However, parental behaviour clearly changed during the high-intensity noise treatment and is suggestive for a compensatory strategy and this warrants attention to further reveal short- and long-term impact of noisy conditions on individual fitness and population consequences.

## MATERIALS AND METHODS

### Subjects and housing

The zebra finches used in this study originated from the breeding colony at Leiden University. Thirty first-time breeding pairs (900±60 days old) were established from a pool of birds that had participated in a noise avoidance test (Liu et al., 2020). The breeding experiment reported here started 142 days after the noise avoidance test. Birds were caught from single-sex aviaries (L×W×H: 2×2×2 m) and unrelated birds were randomly assigned to pairs and each pair was placed into one of 30 identical breeding cages (1×0.5×0.4 m), in one of two identical breeding rooms (3.65×3.05×3. m, for details see Fig. S1). Every breeding cage had an opening (0.09×0.09 m) to hold a white plastic nest box drawer (0.11×0.09×0.09 m) in the top right corner of the cage.

### Sound stimuli

For the noise exposure during breeding, two 24-h recordings of highway traffic noise were chosen from a pool of field recordings that had been collected for our earlier study using SM1 sound meters (Wildlife Acoustics; details see Liu et al., 2020). Noise exposure studies generally compare traffic noise to no playbacks, but without information on birds’ reactions to novel sounds, any observed effect could also arise from the presence of a novel sound rather than from traffic noise per se. We thus used the results from a previous spatial avoidance test (Liu et al., 2020) to select a behaviourally neutral sound (moderate-intensity, far-distance traffic noise) as control and an aversive experimental treatment sound (high-intensity, near-distance traffic noise).

Briefly (details in Liu et al., 2020), the ‘near-distance’ and ‘far-distance’ recordings had been recorded at 15 m (52.098504N, 4.439159E) and 300 m distance (52.103469N, 4.441135E) from the same highway (A4, between Amsterdam and Rotterdam in the Netherlands) during the same 24 h interval. At both locations the sound meters had been placed in open landscape at a height of ca. 0.8 m above the ground such that the microphone pointed across the open landscape towards the highway. Absolute sound pressure levels (SPL) of the first 3 min of each recording in the field were measured by a sound pressure meter (Pulsar Instruments Plc, Model 30, A weighted, reading LAT with an interval of 10 s) and then used later as reference to calculate the average sound levels per 30 s for the entire recordings (see details in Liu et al., 2020): Mean of all 30 s blocks near-distance: 68.7±3.2 dB(A); far-distance: 52.8±5.2 dB(A) re:20 μPa. High amplitudes (>2 standard deviations) were inspected and deleted to avoid startle responses from the birds. For these recordings (and from another location) 30-min-long sequences (extracted from recordings between 11 and 12 am) had been used in the noise avoidance behaviour tests (Liu et al., 2020) that involved the same birds now recruited as parents in the current study. Note that in these previous tests, the parental birds had not changed their behaviour or moved away from the far-distance (moderate level) highway noise, henceforth we selected and designated this moderate-intensity as behaviourally neutral control sound. In contrast, the high-intensity near-distance stimuli had been actively avoided and were thus selected as behaviourally aversive level for the current breeding experiment.

### Traffic noise playbacks

In each breeding room, two loudspeakers (CB4500, Blaupunkt, Hildesheim, Germany) were positioned opposite the breeding cages (details see Fig. S1) and playback levels were adjusted such that the sound level during playback was roughly the same in every breeding cage [aversive noise: cages 70.2±0.5 dB(A), empty nest boxes 68.5±0.9 dB(A); control noise: cages 51.5±0.4 dB(A), empty nest boxes 51.4±0.6 dB(A)], and equivalent to the levels at the recording location when measured with the same sound pressure meter. Because nesting material might act as an acoustic insulator to ambient noise (Potvin, 2019), we also checked after the experiments whether sound levels were attenuated inside the nest boxes when they contained zebra finch nests. To do so, a set of the same type of nest boxes (*N*=18) with completed zebra finch nests inside was obtained from our regular laboratory breeding colony after chicks had left the nests. These nest boxes were then placed into the exact same cages used in this study. We found no measurable differences in sound levels inside these nest boxes with nesting material compared to what we had measured before with empty nest boxes [aversive noise: 69.1±1.1 dB(A), control noise: 52.1±0.4 dB(A)].

### Experimental breeding

All breeding pairs participated in two rounds of breeding while exposed to continuous traffic noise playback of the looped 24 h recordings of either the aversive or the control traffic noise (see Fig. 1). The pairs were moved into the breeding cages 3 days before the onset of the noise playback. From day 3 onwards, the playback faded in from zero to the maximum amplitude level in 14 days until the noise level inside the breeding cages had reached the same level as at the original recording sites. After another week, all pairs were provided with a nest box, hay and coconut fibre to build nests.

All nest boxes were checked daily by one of the experimenters (Q.L., E.G. or K.F.) to track the dates of egg laying and hatching. On the day a chick hatched, it was weighed and individually marked by cutting some of its down feathers (Adam et al., 2014). To break correlations between parental quality, brood size and offspring quality, all broods were fully or partly cross-fostered (within the same treatment room) at age 3.6±1.7 days. Cross-fostered broods were mostly small (2–3 chicks) or large broods (5–6 chicks) to test whether chicks from larger brood sizes that are known to be of lower condition (de Kogel, 1997; Naguib et al., 2004) were more vulnerable to chronic traffic noise exposure. Brood sizes before and after cross-fostering were not correlated (Spearman, r_s_=0.21, *N*=43 broods, *P*=0.18, pairs without offspring were excluded from this analysis). The age composition after the cross-fostering was ideally no more than a day between subsequent hatching dates, but because of asynchronous breeding this was up to 2 days in some nests resulting in a range of 0−2 days and a mean of 1.1±0.2 days.

Chicks were ringed at a median brood age of 11 days, and at 65 days of age, chicks were moved to participate in a noise avoidance test as a first of several behavioural tests (see Fig. 1, Liu et al. in prep) and then moved to single-sex aviaries (2×1.5×2 m, 12−20 birds) in bird stock rooms without playbacks [ambient sound level without birds<40 dB(A)]. The parents remained in the breeding cages while playback in both experimental rooms faded out from the maximum amplitude level to zero over the course of the next week until the previous laboratory ambient level was reached again. The breeding pairs were then given a 2-week playback-free break before the noise playback procedure and a second breeding cycle as described above started again. For this second breeding round, pairs remained in the same cages, but the type of noise playback (aversive or neutral noise) was reversed between rooms (for a full timeline see Fig. 1).

### Breeding outcomes

All breeding cages were checked daily until all chicks had fledged or until more than 40 days since the introduction of the nest boxes had passed without any chicks hatching. For all pairs, we measured the latency to the first egg (in days, counting from the day the nest boxes were provided), the clutch size (as number of eggs laid without a break longer than 3 days), and the number of successfully hatched chicks per pair per breeding round. In the first round, we weighed the eggs of a clutch when females stopped laying eggs for 3 days, but due to an unfortunate combination of personal and technical circumstances, in the second round the clutch weight was not assessed completely and reliably. However, in the second round, we had additionally weighed the chicks on their first day (‘hatchling weight’) when handling them for cutting their down feathers. While this measure will show some variation with respect to the number of hours since hatching, the measure should still detect systematic differences at the group level, and we therefore report this measure to compare reproductive output among the groups in the second round before cross-fostering started. In addition, in both rounds, individual offspring weight was measured when chicks were 11, 65 and 120 days old. On days with video recording, the video recording always preceded the weighing. At 65 days, individual juvenile birds were weighed before participating in a noise avoidance test. After the test, birds were housed with other same-age, same-sex young birds (*N*=30-40) in group aviaries (1/2×2×2 m) without noise playbacks. At 120 days, birds were weighed before participating in a second noise avoidance test.

### Video recording of parental behaviour

During the first breeding round, when the pairs had hatchlings and the median age of a brood reached 5, 8, 11 or 14 days, a camera (Panasonic HC-v500, Osaka, Japan/ JVC, Everio HDD, Kanagawa, Japan/ JVC quad, Kanagawa, Japan) was placed in the middle of the room to simultaneously film all focal cages from the front. These videos were later used to score nest attendance of the parents. To validate whether nest attendance predicted feeding events, in-nest cameras (GoPro HERO 3+, GoPro, CA, USA) that could be fixed under the roof of the nest box were used to record inside the nest. To prevent a neophobia response to the in-nest cameras during filming, dummy cameras (a black cardboard dummy in size and shape resembling the GoPro HERO 3+) had been attached to all nest boxes from day 1 onwards. These were replaced with real cameras only on the day of filming and returned afterwards. Filming always took place between 9:00 to 12:00 am for the duration of 55 min and the schedule was balanced by nest, brood age and treatment. For the second breeding round, when nest attendance had been validated to predict feeding events (see below), only nest attendance was assessed (filming the breeding pairs with chicks with the room camera).

### Video scoring

For all video recordings, the first 5 min were excluded from analyses to make sure birds had enough time to resume normal activities after the camera was installed. Recordings were then analysed with the observers blinded to the treatment from the start of the 6th to the end of the 55th minute to have equally long sequences of 50 min for all recordings. Recordings were scored in BORIS video analysis software (Friard and Gamba, 2016, v.6.1.6). From the room camera recordings, we measured the duration of individual parent’s nest visits by marking the time of all instances from when a bird entered the nest box (=both legs inside) to when the bird left the nest box (=both legs outside). Male and female parents could be told apart by their differences in plumage. The cumulative time each parent spent in the nest box was then obtained by adding up the duration of all the nest visits per recording (‘cumulative nest attendance’). We also calculated a pair’s ‘combined nest attendance’ during the recorded period by adding up the total time each parent spent in the nest box. If both parents stayed in the nest at the same time, both times were added to the count, but this was rare. In only two recordings, parents were seen to stay together in the nest for longer than 5% of the observed time.

In addition, we scored individual feeding events from the recordings of the in-nest cameras used in the first breeding round. Each instance where a parent inserted at least part of its beak into a chick’s open beak gape was counted as a feeding event. Videos were scored at least by one experimenter (E.G.) blinded with respect to the treatments. We firstly established inter- and intra-experimenter fidelity in scoring: two observers (Q.L. and E.G.) separately scored the same five videos and their scores were highly similar (intraclass correlation coefficient=0.99). E.G. also checked her own intra-observer repeatability by rescoring the first three videos after having finished all videos. The initial and later scores of these videos were highly consistent (intraclass correlation coefficient=0.99).

### Statistical analyses

#### Reproductive parameters

To test if noise exposure affected reproductive investment, we created generalised mixed linear models in R (package glmmTMB in R 3.5.3) with the following response variables: (1) latency to the first egg; (2) clutch size; (3) successfully hatched chicks; (4) the number of unhatched eggs. The playback type (aversive/neutral), breeding round (1 or 2, ordered categorical) and their interaction were fixed effects. Biological parent IDs were random intercepts. Effects of the two noise treatments on clutch weight (round 1) and hatchling weight (round 2) were tested with linear models with playback type as fixed effect. For each measurement, we created two models, first including all pairs and then repeated the analyses excluding the pairs that had not laid any eggs. To test if the 11, 65 and 120 days old offspring weights were affected by noise treatment, we created a linear mixed model with offspring weight as the response variable. Playback type and age of the offspring (11/65/120) were both set as categorical variables (because three time points are not sufficient for modelling linearity) and breeding round was fixed effect. Brood size (1–6) was a covariate. We also included the interactions among playback type, age and brood size in this model. Bird IDs and social parent IDs were treated as random intercepts.

As single studies with relatively small sample sizes might produce spurious results (Cooper et al., 2019), we also performed a meta-analysis combining our results on hatching failure (unhatched eggs or embryo mortality) and offspring mass with the data from two other similar studies in which zebra finches were also raised under chronic anthropogenic noise exposure (Potvin and MacDougall-Shackleton, 2015; Zollinger et al., 2020). These two and our study measured mostly the same parameters (hatching success, fledging success, chick mass at different stages) but treatment effects on hatching failure differed between studies, and was either the total number of unhatched eggs (our study, but not reported by other studies) or embryo mortality (Potvin and MacDougall-Shackleton, 2015 and Zollinger et al., 2020). The effect size of noise on offspring mass was calculated by fitting three linear mixed model models of offspring mass (same variables as the overall offspring mass model except for the age of the offspring), one model per age. To calculate the effect size of these models, ‘effectsize’ function (method=‘refit’, robust=TRUE) from the R package ‘effectsize’ (version 0.4.1, Ben-Shachar et al., 2020) was used. We then combined the effect sizes from our and the two aforementioned studies. Since both studies used a Bayesian framework, we directly took their reported ‘mean effect’ for embryo mortality and offspring mass as treatment effect and calculated the accompanying standard error using the following formula: (upper limit of CI – lower limit of CI)/3.92. With the effect sizes from three studies obtained, a meta-analysis was performed on hatching failure and offspring mass: first for each development stage (hatchlings, fledglings and subadults and adults), then combining all development stages. We used the function ‘metagen’ (random-effects model, Hedges’ g, other settings were default) from the R package ‘meta’ (version 4.15-1, Schwarzer, 2007; Balduzzi et al., 2019) to perform all meta-analyses.

#### Parental behaviour

We tested if the sex of the parent affected the nest attendance using a linear mixed model. Nest attendance was the response variable, the sex of the parent (female/male), breeding round (first/second) and the playback type (aversive/control) fixed effects, brood size (1-6) and brood median age (5, 8, 11, 14) covariates. We also added three two-way interactions between the playback type and (1) sex, (2) brood size, (3) brood median age to the model. Social parent IDs were treated as random intercepts. This analysis was repeated, using the same approach and the same factors (without the factor parental sex) for the combined nest attendance.

To test if combined nest attendance predicted feeding events, we created another generalised linear mixed model using data from the first breeding round where we had measured both parental attendance and had scored parental feeding activity using the in-nest videos. The total of the feeding events was the response variable, the scaled (mean=0, s.d.=1 after scaling) combined nest attendance the explanatory variable, the noise playback a fixed effect, brood size and brood median age covariates and social parent IDs random intercepts. We also included the interactions between playback type and (1) brood size and (2) brood median age in this model.

#### Model selection

For all models, whenever an interaction term was not significant, we then compared the full model with a novel model keeping all previous factors but excluding the interaction term(s). If the novel model was significantly better in explaining the data, we treated the novel model as the best model. The report of the models with rejected terms can be found in supplementary data (Tables S2–S4).

#### Ethical note

The experiments described here were reviewed and approved by the committee for animal experimentation at Leiden University and the Centrale Commissie Dierproeven (CCD) of the Netherlands and monitored by the Animal Welfare Body of Leiden University, in accordance with national and European legislation.

## Acknowledgements

We thank our animal caretakers Peter Snelderwaard and Michelle Geers for helping with animal care during the experiments and two anonymous referees and the editor for constructive comments on the manuscript.

## Author contributions

Conceptualization: Q.L., H.S., K.R.; Methodology: Q.L., K.R.; Software: Q.L.; Validation: Q.L.; Formal analysis: Q.L., E.G., K.F.; Investigation: Q.L., E.G., K.F.; Resources: K.R.; Data curation: Q.L.; Writing - original draft: Q.L.; Writing - review & editing: Q.L., H.S., K.R.; Visualization: Q.L.; Supervision: H.S., K.R.; Project administration: H.S., K.R.; Funding acquisition: Q.L, H.S., K.R.

## References

Adam, I., Scharff, C. and Honarmand, M. (2014). Who is who? Non-invasive methods to individually sex and mark altricial chicks. J. Vis. Exp. 51429.

Angelier, F., Meilleŕe, A., Grace, J. K., Trouvé, C. and Brischoux, F. (2016). No evidence for an effect of traffic noise on the development of the corticosterone stress response in an urban exploiter. Gen. Comp. Endocrinol. 232, 43–50. doi:10.1016/j.ygcen.2015.12.007

Arévalo, J. E. and Newhard, K. (2011). Traffic noise affects forest bird species in a protected tropical forest. Rev. Biol. Trop. 59, 969–980.

Balduzzi, S., Rücker, G. and Schwarzer, G. (2019). How to perform a meta-analysis with R: A practical tutorial. Evid. Based. Ment. Health 22, 153–160. doi:10.1136/ebmental-2019-300117

Bayne, E. M., Habib, L. and Boutin, S. (2008). Impacts of chronic anthropogenic noise from energy-sector activity on abundance of songbirds in the boreal forest. Conserv. Biol. 22, 1186–1193. doi:10.1111/j.1523-1739.2008.00973.x

Ben-Shachar, M., Lüdecke, D. and Makowski, D. (2020). effectsize: Estimation of effect size indices and standardized parameters. J. Open Source Softw. 5, 2815. doi:10.21105/joss.02815

Benítez-López, A., Alkemade, R. and Verweij, P. A. (2010). The impacts of roads and other infrastructure on mammal and bird populations: A meta-analysis. Biol. Conserv. 143, 1307–1316. doi:10.1016/j.biocon.2010.02.009

Bennett, V. J. (2017). Effects of road density and pattern on the conservation of species and biodiversity. Curr. Landsc. Ecol. Reports 2, 1–11. doi:10.1007/s40823-017-0020-6

Boucaud, I. C. A., Mariette, M. M., Villain, A. S. and Vignal, C. (2016). Vocal negotiation over parental care? Acoustic communication at the nest predicts partners’ incubation share. Biol. J. Linn. Soc. 117, 322–336. doi:10.1111/bij.12705

Boucaud, I. C. A., Perez, E. C., Ramos, L. S., Griffith, S. C. and Vignal, C. (2017). Acoustic communication in zebra finches signals when mates will take turns with parental duties. Behav. Ecol. 28, 645–656. doi:10.1093/beheco/arw189

Brandl, H. B., Griffith, S. C. and Schuett, W. (2019). Wild zebra finches choose neighbours for synchronized breeding. Anim. Behav. 151, 21–28. doi:10.1016/j.anbehav.2019.03.002

Briga, M., Koetsier, E., Boonekamp, J. J., Jimeno, B. and Verhulst, S. (2017). Food availability affects adult survival trajectories depending on early developmental conditions. Proc. Royal Soc. B 284, 20162287. doi:10.1098/rspb.2016.2287

Brischoux, F., Meilleŕe, A., Dupoué, A., Lourdais, O. and Angelier, F. (2017). Traffic noise decreases nestlings’ metabolic rates in an urban exploiter. J. Avian Biol. 48, 905–909. doi:10.1111/jav.01139

Brumm, H., Goymann, W., Derégnaucourt, S., Geberzahn, N. and Zollinger, S. A. (2021). Traffic noise disrupts vocal development and suppresses immune function. Sci. Adv. 7, eabe2405. doi:10.1126/sciadv.abe2405

Buchanan, K. L., Leitner, S., Spencer, K. A., Goldsmith, A. R. and Catchpole, C. K. (2004). Developmental stress selectively affects the song control nucleus HVC in the zebra finch. Proc. Royal Soc. B 271, 2381–2386. doi:10.1098/rspb.2004.2874

Bunkley, J. P., McClure, C. J. W., Kawahara, A. Y., Francis, C. D. and Barber, J. R. (2017). Anthropogenic noise changes arthropod abundances. Ecol. Evol. 7, 2977-2985. doi:10.1002/ece3.2698

Cooper, H., Hedges, L. V. and Valentine, J. C. (2019). The Handbook of Research Synthesis and Meta-Analysis, 3rd edn. New York, United States: Russell Sage Foundation.

Corbani, T. L., Martin, J. E. and Healy, S. D. (2021). The impact of acute loud noise on the behavior of laboratory birds. Front. Vet. Sci. 7, 607632. doi:10.3389/fvets.2020.607632

Crino, O. L., Van Oorschot, B. K., Johnson, E. E., Malisch, J. L. and Breuner, C. W. (2011). Proximity to a high traffic road: Glucocorticoid and life history consequences for nestling white-crowned sparrows. Gen. Comp. Endocrinol. 173, 323–332. doi:10.1016/j.ygcen.2011.06.001

Crino, O. L., Johnson, E. E., Blickley, J. L., Patricelli, G. L. and Breuner, C. W. (2013). Effects of experimentally elevated traffic noise on nestling white-crowned sparrow stress physiology, immune function and life history. J. Exp. Biol. 216, 2055–2062.

de Kogel, C. H. (1997). Long-term effects of brood size manipulation on morphological development and sex-specific mortality of offspring. J. Anim. Ecol. 66, 167. doi:10.2307/6019

Dorado-Correa, A. M., Zollinger, S. A., Heidinger, B. and Brumm, H. (2018). Timing matters: traffic noise accelerates telomere loss rate differently across developmental stages. Front. Zool. 15, 29. doi:10.1186/s12983-018-0275-8

Evans, J. C., Dall, S. R. X. and Kight, C. R. (2018). Effects of ambient noise on zebra finch vigilance and foraging efficiency. PLoS One 13, e0209471.

Ferraro, D. M., Le, M.-L. T. and Francis, C. D. (2020). Combined effect of anthropogenic noise and artificial night lighting negatively affect Western bluebird chick development. Condor 122, 1–12. doi:10.1093/condor/duaa037

Flores, R., Penna, M., Wingfield, J. C., Cuevas, E., Vásquez, R. A. and Quirici, V. (2020). Effects of traffic noise exposure on corticosterone, glutathione and tonic immobility in chicks of a precocial bird. Conserv. Physiol. 7, coz061. doi:10.1093/conphys/coz061

Francis, C. D., Ortega, C. P. and Cruz, A. (2009). Noise pollution changes avian communities and species interactions. Curr. Biol. 19, 1415–1419.

Francis, C. D., Paritsis, J., Ortega, C. P. and Cruz, A. (2011). Landscape patterns of avian habitat use and nest success are affected by chronic gas well compressor noise. Landsc. Ecol. 26, 1269–1280.

Friard, O. and Gamba, M. (2016). BORIS: a free, versatile open-source event-logging software for video/audio coding and live observations. Methods Ecol. Evol. 7, 1325–1330. doi:10.1111/2041-210X.12584

Griffith, S. C. and Buchanan, K. L. (2010). The zebra finch: the ultimate Australian supermodel. Emu 110, 5–12. doi:10.1071/MUv110n3_ED

Griffith, S. C., Ton, R., Hurley, L. L., McDiarmid, C. S. and Pacheco-Fuentes, H. (2021). The ecology of the zebra finch makes it a great laboratory model but an outlier amongst passerine birds. Birds 2, 60–76. doi:10.3390/birds2010004

Grunst, M. L., Grunst, A. S., Pinxten, R. and Eens, M. (2020). Anthropogenic noise is associated with telomere length and carotenoid-based coloration in free-living nestling songbirds. Environ. Pollut. 260, 114032. doi:10.1016/j.envpol.2020.114032

Grunst, M. L., Grunst, A. S., Pinxten, R. and Eens, M. (2021). Little parental response to anthropogenic noise in an urban songbird, but evidence for individual differences in sensitivity. Sci. Total Environ. 769, 144554. doi:10.1016/j.scitotenv.2020.144554

Habib, L., Bayne, E. M. and Boutin, S. (2007). Chronic industrial noise affects pairing success and age structure of ovenbirds Seiurus aurocapilla. J. Appl. Ecol. 44, 176–184. doi:10.1111/j.1365-2664.2006.01234.x

Halfwerk, W., Holleman, L. J. M., Lessells, C. K. M. and Slabbekoorn, H. (2011a). Negative impact of traffic noise on avian reproductive success. J. Appl. Ecol. 48, 210–219. doi:10.1111/j.1365-2664.2010.01914.x

Halfwerk, W., Bot, S., Buikx, J., van der Velde, M., Komdeur, J., ten Cate, C. and Slabbekoorn, H. (2011b). Low-frequency songs lose their potency in noisy urban conditions. Proc. Natl. Acad. Sci. 108, 14549–14554. doi:10.1073/pnas.1109091108

Halfwerk, W., Both, C. and Slabbekoorn, H. (2016). Noise affects nest-box choice of 2 competing songbird species, but not their reproduction. Behav. Ecol. 27, 1592–1600. doi:10.1093/beheco/arv204

Herrera-Montes, M. I. and Aide, T. M. (2011). Impacts of traffic noise on anuran and bird communities. Urban Ecosyst. 14, 415–427. doi:10.1007/s11252-011-0158-7

Huet des Aunay, G., Grenna, M., Slabbekoorn, H., Nicolas, P., Nagle, L., Leboucher, G., Malacarne, G. and Draganoiu, T. I. (2017). Negative impact of urban noise on sexual receptivity and clutch size in female domestic canaries. Ethology 123, 843–853. doi:10.1111/eth.12659

Injaian, A. S., Taff, C. C., Pearson, K. L., Gin, M. M. Y., Patricelli, G. L. and Vitousek, M. N. (2018a). Effects of experimental chronic traffic noise exposure on adult and nestling corticosterone levels, and nestling body condition in a free-living bird. Horm. Behav. 106, 19–27. doi:10.1016/j.yhbeh.2018.07.012

Injaian, A. S., Poon, L. Y. and Patricelli, G. L. (2018b). Effects of experimental anthropogenic noise on avian settlement patterns and reproductive success. Behav. Ecol. 29, 1181–1189. doi:10.1093/beheco/ary097

Injaian, A. S., Taff, C. C. and Patricelli, G. L. (2018c). Experimental anthropogenic noise impacts avian parental behaviour, nestling growth and nestling oxidative stress. Anim. Behav. 136, 31–39. doi:10.1016/j.anbehav.2017.12.003

Injaian, A. S., Gonzalez-Gomez, P. L., Taff, C. C., Bird, A. K., Ziur, A. D., Patricelli, G. L., Haussmann, M. F. and Wingfield, J. C. (2019). Traffic noise exposure alters nestling physiology and telomere attrition through direct, but not maternal, effects in a free-living bird. Gen. Comp. Endocrinol. 276, 14–21. doi:10.1016/j.ygcen.2019.02.017

Kight, C. R., Saha, M. S. and Swaddle, J. P. (2012). Anthropogenic noise is associated with reductions in the productivity of breeding Eastern bluebirds (Sialia sialis). Ecol. Appl. 22, 1989–1996. doi:10.1890/12-0133.1

Kleist, N. J., Guralnick, R. P., Cruz, A., Lowry, C. A. and Francis, C. D. (2018). Chronic anthropogenic noise disrupts glucocorticoid signaling and has multiple effects on fitness in an avian community. Proc. Natl. Acad. Sci. USA 115, E648–E657. doi:10.1073/pnas.1709200115

Krause, E. T., Krüger, O. and Schielzeth, H. (2017). Long-term effects of early nutrition and environmental matching on developmental and personality traits in zebra finches. Anim. Behav. 128, 103–115. doi:10.1016/j.anbehav.2017.04.003

Kroodsma, D. E. (1976). Reproductive development in a female songbird: differential stimulation by quality of male song. Science 192, 574–575. doi:10.1126/science.192.4239.574

Kunc, H. P. and Schmidt, R. (2019). The effects of anthropogenic noise on animals: a meta-analysis. Biol. Lett. 15, 20190649. doi:10.1098/rsbl.2019.0649

Leonard, M. and Horn, A. (2012). Ambient noise increases missed detections in nestling birds. Biol. Lett. 8, 530–532. doi:10.1098/rsbl.2012.0032

Leonard, M., Horn, A., Oswald, K. and McIntyre, E. (2015). Effect of ambient noise on parent–offspring interactions in tree swallows. Anim. Behav. 109, 1–7. doi:10.1016/j.anbehav.2015.07.036

Lessells, C. M., Riebel, K. and Draganoiu, T. I. (2011). Individual benefits of nestling begging: experimental evidence for an immediate effect, but no evidence for a delayed effect. Biol. Lett. 7, 336–338. doi:10.1098/rsbl.2010.0870

Liu, Q., Slabbekoorn, H. and Riebel, K. (2020). Zebra finches show spatial avoidance of near but not far distance traffic noise. Behaviour 157, 333–362. doi:10.1163/1568539X-bja10004

Liu, Y., Zollinger, S. A. and Brumm, H. (2021). Chronic exposure to urban noise during the vocal learning period does not lead to increased song frequencies in zebra finches. Behav. Ecol. Sociobiol. 75, 1–9. doi:10.1007/s00265-020-02935-9

Loning, H., Griffith, S. C. and Naguib, M. (2021). Zebra finch song is a very short-range signal in the wild: evidence from an integrated approach. Behav. Ecol. 33, 37–46. doi:10.1093/beheco/arab107

Lucass, C., Eens, M. and Müller, W. (2016). When ambient noise impairs parent-offspring communication. Environ. Pollut. 212, 592–597. doi:10.1016/j.envpol.2016.03.015

Mariette, M. M. and Griffith, S. C. (2012). Nest visit synchrony is high and correlates with reproductive success in the wild zebra finch Taeniopygia guttata. J. Avian Biol. 43, 131–140. doi:10.1111/j.1600-048X.2012.05555.x

Mariette, M. M. and Griffith, S. C. (2015). The adaptive significance of provisioning and foraging coordination between breeding partners. Am. Nat. 185, 270–280. doi:10.1086/679441

McClure, C. J. W., Ware, H. E., Carlisle, J., Kaltenecker, G. and Barber, J. R. (2013). An experimental investigation into the effects of traffic noise on distributions of birds: avoiding the phantom road. Proc. Royal Soc. B 280, 20132290–20132290. doi:10.1098/rspb.2013.2290

McIntyre, E., Leonard, M. L. and Horn, A. G. (2014). Ambient noise and parental communication of predation risk in tree swallows, Tachycineta bicolor. Anim. Behav. 87, 85–89. doi:10.1016/j.anbehav.2013.10.013

Meilleŕe, A., Brischoux, F., Angelier, F., Meillere, A., Brischoux, F. and Angelier, F. (2015a). Impact of chronic noise exposure on antipredator behavior: An experiment in breeding house sparrows. Behav. Ecol. 26, 569–577. doi:10.1093/beheco/aru232

Meilleŕe, A., Brischoux, F., Ribout, C. and Angelier, F. (2015b). Traffic noise exposure affects telomere length in nestling house sparrows. Biol. Lett. 11, 20150559. doi:10.1098/rsbl.2015.0559

Mulholland, T. I., Ferraro, D. M., Boland, K. C., Ivey, K. N., Le, M.-L., LaRiccia, C. A., Vigianelli, J. M. and Francis, C. D. (2018). Effects of experimental anthropogenic noise exposure on the reproductive success of secondary cavity nesting birds. Integr. Comp. Biol. 58, 967–976. doi:10.1093/icb/icy079

Naguib, M. (2013). Living in a noisy world: indirect effects of noise on animal communication. Behaviour 150, 1069–1084. doi:10.1163/1568539X-00003058

Naguib, M., Riebel, K., Marzal, A. and Gil, D. (2004). Nestling immunocompetence and testosterone covary with brood size in a songbird. Proc. Royal Soc. B 271, 833–838. doi:10.1098/rspb.2003.2673

Naguib, M., van Oers, K., Braakhuis, A., Griffioen, M., de Goede, P. and Waas, J. R. (2013). Noise annoys: effects of noise on breeding great tits depend on personality but not on noise characteristics. Anim. Behav. 85, 949–956. doi:10.1016/j.anbehav.2013.02.015

Newport, J., Shorthouse, D. J. and Manning, A. D. (2014). The effects of light and noise from urban development on biodiversity: Implications for protected areas in Australia. Ecol. Manag. Restor. 15, 204–214. doi:10.1111/emr.12120

Ng, C. S., Des Brisay, P. G. and Koper, N. (2019). Chestnut-collared longspurs reduce parental care in the presence of conventional oil and gas development and roads. Anim. Behav. 148, 71–80. doi:10.1016/j.anbehav.2018.12.001

Obomsawin, A. P., Mastromonaco, G. F. and Leonard, M. L. (2021). Chronic noise exposure has context-dependent effects on stress physiology in nestling Tree Swallows (Tachycineta bicolor). Gen. Comp. Endocrinol. 311, 113834. doi:10.1016/j.ygcen.2021.113834

Osbrink, A., Meatte, M. A., Tran, A., Herranen, K. K., Meek, L., Murakami-Smith, M., Ito, J., Bhadra, S., Nunnenkamp, C. and Templeton, C. N. (2021). Traffic noise inhibits cognitive performance in a songbird. Proc. Royal Soc. B 288, 20202851.

Pandit, M. M., Eapen, J., Pineda-Sabillon, G., Caulfield, M. E., Moreno, A., Wilhelm, J., Ruyle, J. E., Bridge, E. S. and Proppe, D. S. (2021). Anthropogenic noise alters parental behavior and nestling developmental patterns, but not fledging condition. Behav. Ecol. 32, 747–755. doi:10.1093/beheco/arab015

Patrick, S. C. and Weimerskirch, H. (2014). Personality, foraging and fitness consequences in a long lived seabird. PLoS One 9, e87269. doi:10.1371/journal.pone.0087269

Potvin, D. A. (2019). Mud acts as a noise dampener in Australian passerine nests.Emu -Austral Ornithol. 119, 45–52. doi:10.1080/01584197.2018.1508355

Potvin, D. A. and MacDougall-Shackleton, S. A. (2015). Traffic noise affects embryo mortality and nestling growth rates in captive zebra finches. J. Exp. Zool. Part A Ecol. Genet. Physiol. 323, 722–730. doi:10.1002/jez.1965

Rehling, A., Spiller, I., Krause, E. T., Nager, R. G., Monaghan, P. and Trillmich, F. (2012). Flexibility in the duration of parental care: Zebra finch parents respond to offspring needs. Anim. Behav. 83, 35–39. doi:10.1016/j.anbehav.2011.10.003

Reijnen, R., Foppen, R. and Meeuwsen, H. (1996). The effects of traffic on the density of breeding birds in Dutch agricultural grasslands. Biol. Conserv. 75, 255–260. doi:10.1016/0006-3207(95)00074-7

Santos, E. S. A. and Nakagawa, S. (2012). The costs of parental care: a meta-analysis of the trade-off between parental effort and survival in birds. J. Evol. Biol. 25, 1911–1917. doi:10.1111/j.1420-9101.2012.02569.x

Schroeder, J., Nakagawa, S., Cleasby, I. R. and Burke, T. (2012). Passerine birds breeding under chronic noise experience reduced fitness. PLoS One 7, e39200. doi:10.1371/journal.pone.0039200

Schwarzer, G. (2007). meta: An R package for meta-analysis. R News 7, 40–45.

Senzaki, M., Barber, J. R., Phillips, J. N., Carter, N. H., Cooper, C. B., Ditmer, M. A., Fristrup, K. M., McClure, C. J. W., Mennitt, D. J., Tyrrell, L. P., et al. (2020). Sensory pollutants alter bird phenology and fitness across a continent. Nature 587, 605–609. doi:10.1038/s41586-020-2903-7

Swaddle, J. P. and Page, L. C. (2007). High levels of environmental noise erode pair preferences in zebra finches: implications for noise pollution. Anim. Behav. 74, 363–368. doi:10.1016/j.anbehav.2007.01.004

Villain, A. S., Fernandez, M. S. A., Bouchut, C., Soula, H. A. and Vignal, C. C. (2016). Songbird mates change their call structure and intrapair communication at the nest in response to environmental noise. Anim. Behav. 116, 113–129. doi:10.1016/j.anbehav.2016.03.009

Vitousek, M. N., Jenkins, B. R., Hubbard, J. K., Kaiser, S. A. and Safran, R. J. (2017). An experimental test of the effect of brood size on glucocorticoid responses, parental investment, and offspring phenotype. Gen. Comp. Endocrinol. 247, 97–106. doi:10.1016/j.ygcen.2017.01.021

Waas, J. R., Colgan, P. W. and Boag, P. T. (2005). Playback of colony sound alters the breeding schedule and clutch size in zebra finch (Taeniopygia guttata) colonies. Proc. Royal Soc. B 272, 383–388. doi:10.1098/rspb.2004.2949

Walthers, A. R. and Barber, C. A. (2020). Traffic noise as a potential stressor to offspring of an urban bird, the European starling. J. Ornithol. 161, 459–467. doi:10.1007/s10336-019-01733-z

Williams, T. D. (2018). Physiology, activity and costs of parental care in birds. J. Exp. Biol. 221, jeb169433. doi:10.1242/jeb.169433

Williams, D. P., Avery, J. D., Gabrielson, T. B. and Brittingham, M. C. (2021). Experimental playback of natural gas compressor noise reduces incubation time and hatching success in two secondary cavity-nesting bird species. Condor 123, 1–11. doi:10.1093/ornithapp/duaa066

Zollinger, S. A., Dorado-Correa, A., Goymann, W., Forstmeier, W., Knief, U., BastidasUrrutia, A. M. and Brumm, H. (2020). Traffic noise exposure depresses plasma corticosterone and delays offspring growth in breeding zebra finches. Conserv. Physiol. 7, 20162287. doi:10.1093/conphys/coz056

