## Supplementary data for "An experimental test of chronic traffic noise exposure on parental behaviour and reproduction in zebra finches"

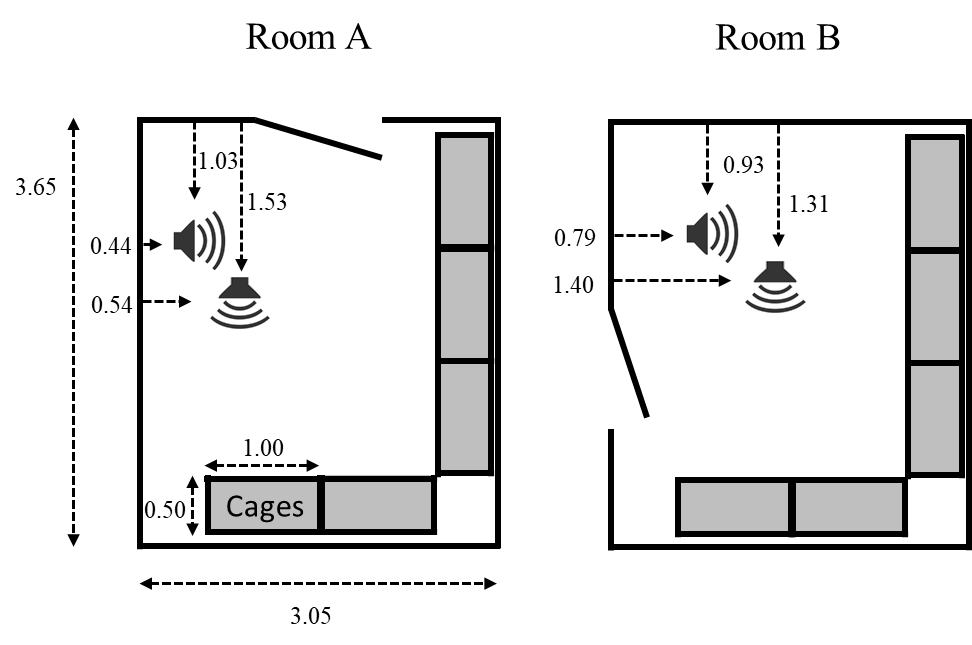

**Figure S1:** **Top view of the two breeding rooms showing the location of the loudspeakers (speaker signs) and breeding cages (grey boxes)**. In each room, blocks of cages were stacked three rows high along two of the walls of the room. The cages in the lowest row were situated 0.6 m above the ground on supporting cabinets (1 x 0.5 x 0.6 m). All measurements are in meters.

**Additional details on experimental procedures and biometric measurements**

*Individual marking of chicks and ringing*

All nestboxes could be opened from outside the cages and were checked daily by one of the experimenters (QL, EG and KF). When a chick hatched, for each chick within a brood the down feathers were cut in an individual specific pattern (head, back, leg or one wing; see Adam et al., 2014 for details) that served as individual ID until the birds were old enough to receive a leg ring. When the median age of a brood reached 11 days, chicks were banded with an orange plastic numbered ID-ring on their left leg.

*Weighing*

Individual offspring weight was measured when individual chicks were 11, 65 and 120 days old. Weight measures were taken as follows: For weighing the very young chicks (at 1 and 11 days), the experimenter prepared a temporary replacement nest with hay and coconut fibres and put this on the weighing dish of a digital balance (Sartorius, BL600, Göttingen, Germany +/- 0.1g). The balance was reset to zero before adding the chick to the dummy nest. To transfer the chick to the dummy nest, the experimenter inserted a partition in the middle of the cage at a moment when both parents were on the side without the nestbox. This created temporarily two compartments, one containing the parents and the other the nestbox. The nestbox could now be swiftly removed from the cage by opening the nestbox drawer. Chicks were identified by their down feather cuts and each chick scheduled for weighing was quickly transferred to the dummy nest on the balance and immediately afterwards returned to its own nestbox. At 65 days, individual juvenile birds were caught from their cages with a net and put in a bag to be weighed on the same balance. At 120 days, individual birds were caught with a net from their aviary and briefly put in a bag to be weighed on the same balance.

*Video recordings*

Before starting a video recording, an experimenter (QL, EG or KF) first carefully approached the cage (avoiding sudden or noisy movements), and then used an opaque plastic divider to divide the cage in two compartments, taking care to have the pair on one side and the nest on the other. Then, the experimenter opened the nestbox drawer, replaced the dummy with a real, already switched on camera, returned the nestbox and removed the inserted partition. Upon leaving the room, the experimenter switched on the room camera to film all focal cages from the front. Fifty-five minutes later, the room camera was switched off and in-nest cameras were replaced with dummy cameras. For the second breeding round, to keep procedures identical to the first breeding round, the experimenter repeated the movements of placing and removing the in-nest cameras at the beginning and at the end of what would have been the recording periods.

| **Table S1**. Breeding outcomes per pair during exposure to either the high-intensity aversive or low-intensity control sound. | | | | | | | | | | | | | | |
| --- | --- | --- | --- | --- | --- | --- | --- | --- | --- | --- | --- | --- | --- | --- |
|  | Latency to the first egg [days]^1^ | | | Clutch size | | | Successfully hatched chicks | | Clutch weight [g]^2^ | | | | Hatchling weight [g]^3^ | |
| **Pair ID** | **A**versive | | **C**ontrol | | **A** | **C** | **A** | **C** | **A** | | **C** | **A** | | **C** |
| **1** | 7 | | 6 | 4 | | 4 | 0 | 0 | 5.5 | na | | na | | - |
| **2** | 6 | | 7 | 6 | | 6 | 5 | 5 | 7.9 | na | | na | | 5.7 |
| **3** | 4 | | 4 | 8 | | 9 | 3 | 7 | 7.2 | na | | na | | 7.2 |
| **4** | 2 | | 7 | 6 | | 4 | 4 | 4 | 5.3 | na | | na | | 3.1 |
| **5** | 2 | | 6 | 5 | | 6 | 1 | 4 | 4.8 | na | | na | | 3.8 |
| **6** | - | | 7 | 0 | | 6 | 0 | 5 | - | na | | na | | 4.7 |
| **7** | 8 | | - | 0 | | 0 | 0 | 0 | 1.3 | na | | na | | - |
| **8** | 2 | | 5 | 7 | | 6 | 1 | 0 | 6.3 | na | | na | | - |
| **9** | - | | 5 | 0 | | 3 | 0 | 1 | - | na | | na | | 0.8 |
| **10** | - | | 10 | 0 | | 3 | 0 | 0 | - | na | | na | | - |
| **11** | 5 | | 3 | 4 | | 4 | 0 | 2 | 4.3 | na | | na | | 1.6 |
| **12** | 6 | | 6 | 5 | | 4 | 0 | 2 | 5.9 | na | | na | | 1.7 |
| **13** | 7 | | 3 | 6 | | 6 | 6 | 5 | 6.4 | na | | na | | 4.7 |
| **14** | 3 | | 3 | 6 | | 5 | 5 | 3 | 5.5 | na | | na | | 2.4 |
| **15** | 3 | | 3 | 4 | | 5 | 1 | 1 | 3.7 | na | | na | | 1.0 |
| **16** | 2 | | 8 | 3 | | 6 | 1 | 6 | na | 4.6 | | 1.3 | | na |
| **17** | 17 | | - | 3 | | 0 | 3 | 0 | na | - | | 2.7 | | na |
| **18** | 4 | | 2 | 5 | | 4 | 0 | 0 | na | 4.2 | | - | | na |
| **19** | 11 | | 2 | 6 | | 4 | 0 | 3 | na | 4.5 | | - | | na |
| **20** | 6 | | 5 | 5 | | 5 | 5 | 2 | na | 6.0 | | 4.8 | | na |
| **21** | 5 | | 2 | 2 | | 3 | 0 | 1 | na | 2.7 | | - | | na |
| **22** | 6 | | 6 | 4 | | 4 | 2 | 3 | na | 5.1 | | 2.2 | | na |
| **23** | - | | 9 | 0 | | 5 | 0 | 0 | na | - | | - | | na |
| **24** | 11 | | 4 | 5 | | 5 | 4 | 0 | na | 5.3 | | 3.7 | | na |
| **25** | - | | - | 0 | | 0 | 0 | 0 | na | - | | - | | na |
| **26** | 5 | | 2 | 5 | | 4 | 3 | 3 | na | 4.7 | | 2.5 | | na |
| **27** | 4 | | 2 | 6 | | 2 | 5 | 0 | na | 1.0 | | 4.7 | | na |
| **28** | 4 | | 7 | 6 | | 6 | 5 | 6 | na | 7.2 | | 4.7 | | na |
| **29** | 2 | | 6 | 11 | | 5 | 1 | 5 | na | 5.4 | | 0.7 | | na |
| **30** | 3 | | 1 | 3 | | 5 | 0 | 0 | na | - | | - | | na |
| **All pairs** | **mean** | - | - | 4.2 | | 4.3 | 1.8 | 2.2 | - | - | | - | | - |
|  | **s.d.** | - | - | 2.4 | | 2.0 | 2.1 | 2.3 | - | - | | - | | - |
| **Bred pairs^4^** | **mean** | 5.4 | 4.9 | 5 | | 4.7 | 2.2 | 2.5 | - | - | | - | | - |
|  | **s.d.** | 3.5 | 2.4 | 2.1 | | 1.4 | 2.1 | 2.3 | - | - | | - | | - |
| *^1^ days were counted after providing nesting materials ^2^ measured in the first round ^3^ measured in the second round* **^4^** *at least one hatched chick* | | | | | | | | | | | | | | |

| **Table S2**: Rejected interaction terms of the generalised linear mixed model of breeding outcomes. The best models are reported in Table 2. | | | |
| --- | --- | --- | --- |
| **Effects** |  | **χ²** | **p-value** |
| a) Latency to the first egg | |  |  |
| Noise |  | 3.86 | 0.06 |
| Round |  | 0.13 | 0.72 |
| Noise * round |  | 0.01 | 0.92 |
| b) Clutch size | |  |  |
| Noise |  | 0.52 | 0.81 |
| Round |  | 1.00 | 0.32 |
| Noise * round |  | 0.16 | 0.69 |
| c) Successfully hatched chicks | |  |  |
| Noise |  | 1.25 | 0.26 |
| Round |  | 1.25 | 0.26 |
| Noise * round |  | 00.01 | 0.92 |
| d) Number of unhatched eggs | |  |  |
| Noise |  | 0.62 | 0.43 |
| Round |  | 0.08 | 0.78 |
| Noise * round |  | 0.02 | 0.88 |
| *Marginal and conditional R^2^ for the models are a) 0.04 and 0.06, b) 0.01 and 0.35, c) 0.02 and0.65, d) 0.01 and 0.43. Biological parents’ IDs were treated as random intercepts.* | | | |

| **Table S3**: Rejected interaction terms of the linear mixed model of offspring weight measured at 11, 65 and 120 days old. | | | | | | | |
| --- | --- | --- | --- | --- | --- | --- | --- |
| **Effects** |  | **Estimate** | **Std.Error** | | | **χ²** | **P value** |
| Noise |  |  | |  | 0.41 | | 0.52 |
| **Age** |  |  | |  | **1544.31** | | **<0.001** |
| Round |  |  | |  | 3.45 | | 0.06 |
| **Brood size** | *Covariate* | *-0.60* | | *0.23* | **11.85** | | **< 0.001** |
| Noise * age | |  | |  | 1.31 | | 0.52 |
| Noise * brood size |  |  | |  | 1.70 | | 0.19 |
| Noise * age * brood size | |  | |  | 1.13 | | 0.58 |
| *Marginal and conditional R^2^ for the model are 0.70 and 0.84. Social parents’ IDs and bird ID were treated as random intercepts.* | | | | | | | |

| **Table S4**: Nest attendance of the parents during the different noise exposures. The table reports the rejected interaction terms of the generalised linear mixed model analysis. The best model is reported in Table 3. | | | | | | | | | | |
| --- | --- | --- | --- | --- | --- | --- | --- | --- | --- | --- |
| **Effects** |  | | | | **Estimate** | **Std.Error** | | | **χ²** | **P value** |
| a) Nest attendance per parent | | | | | | | | | | |
| **Sex** |  | | | |  | |  | **11.11** | | **< 0.001** |
| **Noise** |  | | | |  | |  | **11.00** | | **< 0.001** |
| Round |  | | | |  | |  | 0.21 | | 0.65 |
| **Brood size** | *Covariate* | | | | *-0.07* | | *0.01* | **54.10** | | **< 0.001** |
| **Brood median age** | | | *Covariate* | | *-0.04* | | *0.01* | **83.29** | | **< 0.001** |
| Noise * sex |  | | | |  | |  | 0.39 | | 0.53 |
| Noise * brood size | | | | |  | |  | 0.92 | | 0.34 |
| Noise * brood median age | | | | |  | |  | 0.40 | | 0.53 |
| b) Combined nest attendance | | | | | | | | | | |
| **Noise** |  | | | |  | |  | **13.46** | | **< 0.001** |
| Round |  | | | |  | |  | 0.15 | | 0.70 |
| **Brood size** | *Covariate* | | | | *-0.07* | | *0.001* | **55.48** | | **< 0.001** |
| **Brood median age** | | *Covariate* | | | *-0.03* | | *0.00* | **104.15** | | **< 0.001** |
| Noise * brood size |  | | | |  | |  | 0.97 | | 0.32 |
| Noise * brood median age | | | | |  | |  | 0.50 | | 0.48 |
| c) Combined nest visits | | | | | | | | | | |
| Noise |  | | | |  | |  | 0.33 | | 0.56 |
| **Round** |  | | | |  | |  | **9.66** | | **0.002** |
| Brood size | *Covariate* | | | | -0.12 | | 0.07 | 1.67 | | 0.20 |
| Brood median age | *Covariate* | | | | 0.00 | | 0.02 | 0.02 | | 0.88 |
| Noise * brood size |  | | | |  | |  | 0.06 | | 0.80 |
| Noise * brood median age | | | | |  | |  | 1.19 | | 0.27 |
| d) Feeding events (only measured in the first round) | | | | | | | | | | |
| **Nest attendance** | | *Covariate* | | | *0.07* | | *0.03* | **4.56** | | **0.03** |
| Noise |  | | | |  | |  | 1.50 | | 0.22 |
| Brood size |  | | | | *0.56* | | *0.13* | 15.80 | | <0.001 |
| **Brood median age** | | | *Covariate* | | *-0.08* | | *0.01* | **116.00** | | **< 0.001** |
| **Noise * brood size** | | | | |  | |  | **6.02** | | **0.01** |
| Noise * brood median age | | | | |  | |  | 0.00 | | 0.99 |
| e) Feeding events for brood median age 5 | | | | | | |  |  | |  |
| **Nest attendance** | | | | *Covariate* | *0.17* | | *0.05* |  | | **0.001** |
| **Noise** | | | |  | *0.86* | | *0.16* |  | | **< 0.001** |
| **Brood size** | | | | *Covariate* | *0.33* | | *0.03* |  | | **< 0.001** |
| **Noise * brood size** | | | | | *-0.16* | | *0.04* |  | | **< 0.001** |
| *Marginal and conditional R2 for the models are a) 0.36 and 0.36, b) 0.56 and 0.58, c) 0.09 and 0.11, d) 0.61 and 0.98. Social parents’ IDs were treated as random intercepts* | | | | | | | | | | |

| **Table S5**: The number of successfully hatched chicks of this experiment and of a normal round of breeding, analysed in a generalised linear mixed effect model. | | | | |
| --- | --- | --- | --- | --- |
| **Effects** |  |  | **χ²** | **p-value** |
| Rearing condition | |  | 1.61 | 0.45 |
| *Marginal and conditional R^2^ for this models are 0.01 and 0.68. Biological parents’ IDs were treated as random intercepts.* | | | | |

| **Table S6**: Reproductive output in relation to noise treatment. Linear model analysis with response variable clutch weight (1^st^ round) , total hatchling weight (2^nd^ round). | | | | | | | |
| --- | --- | --- | --- | --- | --- | --- | --- |
| **Effects** |  | **Estimate** | **Std.Error** | | | **t** | **P value** |
| Clutch weight in the 1^st^ breeding round | | | | | | | |
| Noise: aversive | *with non-breeding* | *0.11* | | *0.92* | 0.12 | | 0.93 |
|  | *without* | *1.01* | | *0.57* | 1.75 | | 0.09 |
| Total hatching weight in the 2^nd^ breeding round | | | | | | | |
| Noise: aversive | *with non-breeding* | *0.63* | | *0.78* | 0.81 | | 0.43 |
|  | *without* | *0.52* | | *0.82* | 0.63 | | 0.53 |
